# From Sparse Visits to Continuous Molecular Trajectories with Flow Matching

**DOI:** 10.64898/2026.10.02.756304

**Authors:** Peyman Passban, Shikhar Gupta, Bryan Guan, Tanya Roosta

**Author notes:** This work was conducted independently and is not associated with the author’s employer. All views and conclusions are solely those of the author.

## Abstract

Longitudinal transcriptomic studies often capture only a few molecular snapshots of each patient over time. This makes it challenging to understand how patients are progressing, compare them when progression rates differ, and capture molecular changes between observed visits. We introduce CohortFM,^2^ a flow-matching framework that reconstructs continuous patient-specific molecular trajectories from sparse longitudinal observations and organizes them through an integrated mixture model into a compact set of cohort-level progression patterns. CohortFM separates molecular progression from calendar time, places patients on a shared progression scale, uses observed intermediate states to guide the learned trajectories, and maps the resulting patterns back to real patient samples for biological interpretation. We evaluated CohortFM across COVID-19, sepsis, influenza, and tuberculosis. Despite different sampling patterns, the method generalizes well and consistently produces meaningful trajectories. These trajectories capture biological differences between patients that cannot be explained by the observed samples alone. Compared with state-of-the-art trajectory baselines, CohortFM produces more biologically distinct trajectories and a more complete representation of the patient population. Its finer progression scale also reveals biological programs and temporal dynamics that are not captured by visit-level analysis.

We further evaluated interpolation and true forward prediction to test whether CohortFM can accurately recover unobserved states. CohortFM remains competitive with strong interpolation baselines and is among the strongest learned predictors in the harder forecasting setting. Ablation experiments further show that flow matching is the core learning mechanism and that our design choices improve continuous trajectory learning. Our results support CohortFM as a practical framework for turning sparse molecular snapshots into continuous, biologically interpretable trajectories. By going beyond visit labels, CohortFM makes it possible to study disease progression and predict future molecular states even when patients are sampled only a few times^3^.

## 1 Introduction

Longitudinal molecular profiling allows disease to be studied as a dynamic process rather than as a collection of isolated biological states. Repeated measurements from the same patient can reveal how molecular programs change during disease progression or treatment. In practice, however, longitudinal transcriptomic studies often contain only a handful of measurements per patient. These measurements are high-dimensional and noisy, are collected at irregular intervals, and may be separated by long unobserved periods [Wei et al., 2025, Taheriyoun et al., 2026]. For example, in the tuberculosis dataset used in our experiments, samples are collected at Weeks 0, 1, 4, and then not again until Week 24. With so few observations, the trajectory followed by an individual patient is fundamentally underdetermined. Existing approaches therefore often rely on information from other patients or from the cohort as a whole [Sun et al., 2017, Signorelli et al., 2021, Şapcı et al., 2023]. While this provides statistical support, it can also blur patient-specific behavior.

A related challenge is that *calendar time* is not equivalent to molecular progression. Patients sampled at the same nominal visit may occupy different molecular states, while patients sampled at different times may be at similar stages of progression [Trapnell et al., 2014, Qing et al., 2026]. Grouping samples only by visit can therefore mix distinct progression states and obscure heterogeneous or transient responses. Understanding disease heterogeneity requires not only identifying molecular states, but also modeling how individual patients move between them.

Several classes of methods address parts of these challenges. Time-series clustering identifies recurrent expression patterns [Ernst and Bar-Joseph, 2006]; pseudotime methods infer latent biological orderings; and continuous-time neural models represent irregularly sampled dynamics [Chen et al., 2018, Rubanova et al., 2019]. More recently, methods such as Truffle [Hasanaj et al., 2024] have targeted sparse clinical transcriptomic cohorts by sharing information across patients to construct temporally constrained trajectories. However, sparse clinical molecular data create a harder problem: models need to borrow strength from the cohort without losing what’s specific to each patient, and they need to reconstruct unobserved states without drifting from what was actually measured.

We introduce CohortFM, which approaches this problem by reconstructing continuous patient-specific molecular trajectories rather than limiting the analysis to observed visits. The goal is not simply to generate additional points, but to recover unobserved molecular states accurately enough to preserve patient-specific progression. CohortFM uses flow matching [Lipman et al., 2023] to learn shared dynamics across the cohort while keeping each reconstructed path guided by the patient’s own observed molecular states. It also separates molecular progression from calendar time and places patients on a shared progression scale. The resulting patient trajectories are summarized into a small set of cohort-level progression patterns and mapped back to real patient samples for biological interpretation. The central idea is therefore to use cohort-level information to make sparse trajectory reconstruction possible without sacrificing individual progression.

We evaluate CohortFM across COVID-19, sepsis, severe influenza, and tuberculosis. Across these disease settings, the method produces biologically meaningful trajectories, reveals progression structure not captured by visit-level analysis, and performs competitively on interpolation and strongly on forward prediction. Ablation studies further support flow matching as the core learning mechanism. Our experimental results position CohortFM as a framework for turning sparse and noisy molecular observations into continuous, biologically interpretable patient trajectories.

The remainder of the manuscript is organized to support different levels of engagement with the method. Section 2 reviews related work and the main baseline approaches. Section 3 describes the datasets, their selection, and preprocessing. Section 4 provides a high-level overview of CohortFM, while Section 5 summarizes its main empirical findings. Section 6 then presents the model architecture, objectives, and training procedure in detail, and Section 7 provides the corresponding biological and predictive analyses. The remaining sections discuss broader findings, limitations, and future directions, with additional technical details and supporting experiments provided in the appendix.

## Related Work

Recovering biological progression from high-dimensional molecular measurements has been studied extensively, particularly in single-cell transcriptomics. Pseudo-time methods such as Monocle [Trapnell et al., 2014] and diffusion pseudo-time [Haghverdi et al., 2016] infer latent progression orderings from molecular similarity, while Tempora [Tran and Bader, 2020] and psupertime [Macnair et al., 2022] incorporate temporal information more explicitly. These approaches are effective for organizing molecular states, but sparse clinical cohorts introduce an additional requirement. Repeated measurements from the same patient must be connected while preserving patient-specific progression.

A particularly relevant method is Truffle [Hasanaj et al., 2024], developed for short longitudinal clinical transcriptomic studies. Truffle constructs a graph of observed molecular states and formulates trajectory reconstruction as a multi-commodity-flow problem. Each patient transition is treated as a source–target pair, and the algorithm searches for a low-cost, time-respecting path that may pass through samples from other patients. This allows cohort information to fill gaps in an individual patient’s observed sequence. Truffle is therefore a strong baseline. It directly targets sparse clinical cohorts, preserves temporal constraints, and shares information across patients. Its main distinction from CohortFM is that Truffle reconstructs trajectories through molecular states already observed somewhere in the cohort, whereas CohortFM learns continuous dynamics that can generate unobserved states. Since Truffle has already demonstrated strong performance relative to earlier approaches (such as Tempora, psupertime, and cluster-based methods), we use it as our main trajectory-level baseline.

A central goal of CohortFM is not only to organize observed samples into trajectories, but also to reconstruct missing intermediate states and predict future molecular states. This creates a second evaluation challenge. Generated states must remain accurate despite the small number of observations per patient. We therefore compare CohortFM with strong continuous neural models for temporal molecular prediction. We selected scNODE [Zhang et al., 2024] because it was specifically developed to learn continuous molecular dynamics and predict states at unseen time points. It combines a variational auto-encoder with a neural ODE [Chen et al., 2018] and reports state-of-the-art predictive performance relative to preceding methods across several temporal single-cell datasets. Rather than comparing CohortFM separately with a long list of earlier models, scNODE provides a strong reference that has already demonstrated improvements over them. Importantly, it also uses a different modeling mechanism from CohortFM, allowing us to test whether CohortFM is competitive with a strong continuous-time neural predictor rather than benefiting only from flow matching.

We also selected scFM [Pu et al., 2026], which uses conditional flow matching to model temporal molecular dynamics. It provides a complementary baseline because scFM belongs to the same broad flow-matching family as CohortFM. Comparing the two helps distinguish the benefit of flow matching itself from the specific design choices introduced in CohortFM. Using scNODE and scFM in our comparison helps us address whether CohortFM is competitive with strong neural predictors more generally, and whether its performance can be explained simply by the use of flow matching and our tailored design. The prediction setup makes the task particularly challenging for CohortFM. scNODE and scFM were developed primarily for time-resolved single-cell populations, where observations at different time points are generally unpaired. In our setting, predictions belong to a specific patient trajectory and must therefore be both accurate molecular states and consistent with that patient’s progression. Our evaluation reflects these two requirements. Truffle tests whether CohortFM can recover biologically meaningful and non-redundant progression structure from sparse clinical data, while scNODE and scFM test whether its intermediate and future states are competitive with strong continuous neural predictors.

## 3 Datasets

The primary dataset used for the comprehensive evaluation of our method is the longitudinal COVID-19 transcriptomic cohort (GSE212041),^4^ which profiles temporal changes in neutrophils during SARS-CoV-2 infection using bulk RNA sequencing of enriched peripheral-blood neutrophils. The structure of this dataset is representative of the longitudinal transcriptomic datasets commonly used in similar studies. In general, such datasets consist of a gene-expression matrix in which each sample corresponds to a particular patient at a specific visit, with expression measured across thousands of genes. The number of observations per patient is typically small, often comprising only a handful of visits collected over several days or weeks. Our datasets follow this same general structure.

For the COVID-19 dataset specifically, the full cohort includes 306 SARS-CoV-2-positive patients, 78 symptomatic negative individuals, and 8 healthy controls. Samples were collected primarily at hospital admission (D0), Day 3 (D3), and Day 7 (D7), with additional event-driven measurements (DE) available for a subset of patients. Because longitudinal transcriptomic datasets are typically sparse and noisy, a series of preprocessing steps is required before they can be used for downstream modeling. In this dataset, we retained 650 longitudinal measurements from 304 COVID-19-positive patients (2 out of those 306 were not usable), together with samples from 8 healthy individuals as a reference cohort. We restricted the analysis to protein-coding genes and retained genes expressed above 0.25 CPM in at least 1% of samples. For duplicated gene identifiers, we retained the entry with the highest mean expression. Between-sample differences in sequencing depth and RNA composition were corrected using trimmed mean of M-values normalization [Robinson and Oshlack, 2010], and batch-associated variation was removed using ComBat where batch information was available [Johnson et al., 2007]. Finally, genes were ranked by interquartile range (IQR), and the 2,000 most variable genes were selected for representation learning, yielding a 2,000-dimensional gene-expression vector for each sample. We also evaluated larger feature sets but found that 2,000 genes were sufficient for the downstream analyses.

These preprocessing steps are standard for this type of transcriptomic dataset. We also followed the preprocessing procedure reported by Hasanaj et al. [2024], our main baseline, as closely as possible to ensure a fair comparison with the previously reported results. The resulting 2,000-dimensional expression vector represents the molecular state of a patient at a given visit and serves as the input to the downstream machine-learning and trajectory-modeling pipeline. Thus, the first step of CohortFM begins directly from these 2,000-dimensional sample representations.

As noted earlier, the COVID-19 dataset was selected as our primary case study because it provides a strong baseline for comparison and has a size and sampling structure that meaningfully challenge the method. It is not unusually large, making it representative of the relatively small longitudinal datasets commonly encountered in this field. It also contains only a few scheduled visits, with incomplete follow-up for some patients. Together, these characteristics provide a realistic test of CohortFM under sparse and imperfect sampling. To assess whether the method generalizes beyond this setting, we additionally evaluated CohortFM on three longitudinal transcriptomic datasets covering sepsis, severe influenza, and tuberculosis. Summary statistics for all datasets are provided in Table

The datasets summarized in the table were not chosen at random; they were intentionally selected to cover different longitudinal sampling patterns and challenge CohortFM in complementary ways. Sepsis provides a relatively dense sampling schedule, with daily measurements from Day 1 to Day 5. This allows us to test whether CohortFM can still add value when observations are already closely spaced, for example by reconstructing smoother intermediate progression between visits. Sepsis is also a considerably smaller dataset that makes it even more challenging and interesting for our experiments. Severe influenza is more similar to the COVID-19 setting and provides a useful test of whether the method generalizes to a different disease and cohort with a comparable level of sparsity. Tuberculosis is especially interesting. For this dataset, samples are collected at Weeks 0, 1, and 4, followed by a long gap until Week 24. Since one of the goals of CohortFM is to reconstruct progression across widely separated visits, this dataset provides a particularly useful test of how the model handles long unobserved intervals. Details of the preprocessing and cleaning procedures for these datasets are provided in Appendix A.

## 4 Method Overview

In this section, we describe how CohortFM transforms raw longitudinal molecular data into a small set of cohort-level progression archetypes. Each archetype represents a prototypical molecular trajectory and captures a distinct pattern of progression within the cohort. As noted previously, this section focuses on the conceptual mechanism of the framework, namely how patient-level trajectories are reconstructed and subsequently summarized into cohort-level archetypes. Full technical details are provided in Section 6. The process moves through four successive representations: *molecular measurements→ latent molecular states→ continuous patient trajectories→ cohort-level trajectories*.

The key design principle is to reconstruct each patient’s progression before compressing information across the cohort, rather than clustering samples or patients directly. The input is a longitudinal dataset of several patients, each observed at a small number of clinical visits. At every visit, the patient’s molecular state is represented by a high-dimensional gene-expression vector. Modeling trajectories directly in this space is challenging because of its dimensionality, noise, and sparsity. Each observation is therefore mapped to a compact latent molecular state, that is, a lower-dimensional vector. Each patient is then represented as an ordered sequence of latent vectors, one for each observed visit. This representation is obtained using a variational auto-encoder.

We know that calendar time does not necessarily correspond to molecular progression; for example, two patients sampled after the same number of days may have undergone very different degrees of change. Thus, we represent calendar time and molecular progression separately and assign each patient a normalized molecular-progression coordinate that reflects how far the patient has moved through molecular state space. Conceptually, this is determined by measuring the molecular change between consecutive sampling points and expressing each change relative to the total molecular change observed over that patient’s trajectory.

To make trajectories from different patients directly comparable, each trajectory is mapped to a common normalized progression axis *s*∈ [0, 1]. Every patient is then represented with the same trajectory length and latent dimensionality, even when the original number or timing of visits differs. Accordingly, corresponding positions along this axis represent relative molecular progression rather than equal amounts of elapsed calendar time. Once the patient representations are constructed, a flow-matching neural model, specifically tailored to sparse longitudinal transitions of this form, is used to learn patient-specific trajectories. A patient’s trajectory describes how the molecular state evolves from one observed state to the next. The output of this stage is a continuous latent trajectory anchored by the patient’s observed molecular states.

Finally, CohortFM summarizes the patient-specific trajectories using a mixture of cohort-level trajectories. Rather than assigning every patient rigidly to a single trajectory, the mixture model produces a probability of association between each patient and each cohort-level trajectory, allowing an individual patient trajectory to reflect characteristics of multiple progression patterns. Once learned, these cohort-level trajectories provide the basis for downstream analysis. We examine the biological processes associated with each trajectory, how molecular programs evolve along its progression axis, how the trajectories differ from one another, and how patients are distributed across them. A high-level visualization of the complete CohortFM pipeline is provided in Figure 1.

**Figure 1.**
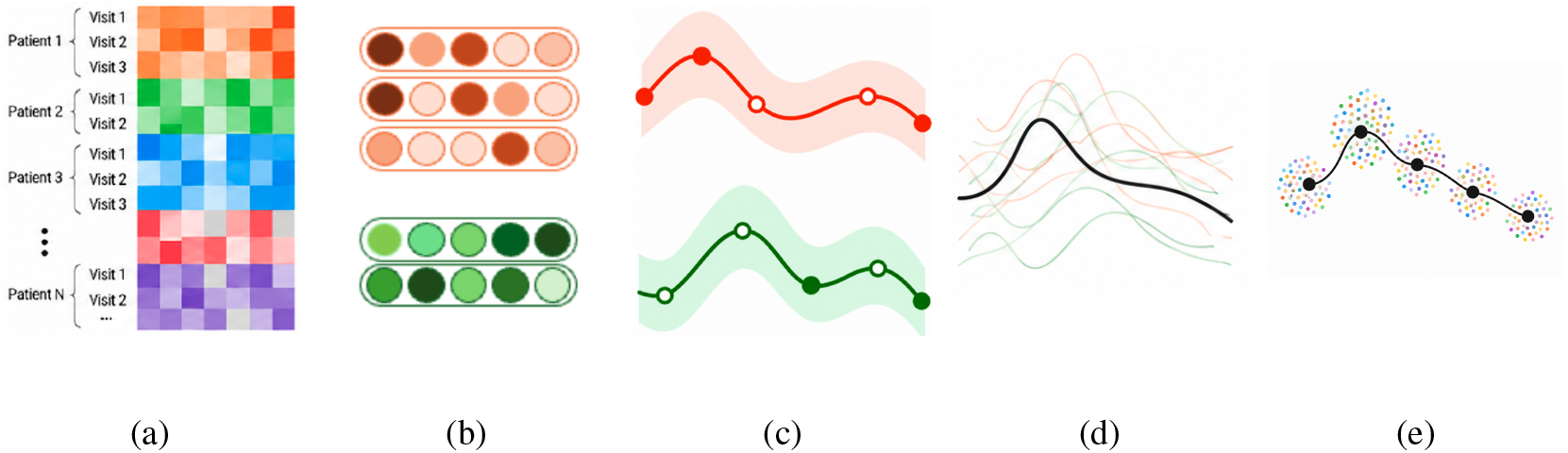
This figure provides a high-level, conceptual overview of the CohortFM framework. (a) A schematic gene-expression matrix in which columns represent genes and rows correspond to samples collected from individual patients at specific time points. (b) Each high-dimensional sample is compressed into a dense, latent embedding. In this illustrative example, one patient has three observed samples and another has two. (c) CohortFM transforms these sparse, discrete observations into continuous patient-specific molecular trajectories. (d) The patient-level trajectories are then combined to illustrate how cohort-level progression patterns can be constructed. This panel is intended only as a conceptual illustration; the actual aggregation procedure is more complex. (e) Finally, the cohort-level trajectory is augmented with patient clouds. At selected positions along the trajectory, patients whose molecular states are close to that position are mapped to the corresponding local region for biological analysis of that state of the trajectory.

## 5 Key Results

We evaluated CohortFM on four longitudinal transcriptomic disease cohorts (COVID-19, sepsis, severe influenza, and tuberculosis) covering substantially different cohort sizes, diseases, and sampling schedules. The evaluation was designed to test three central questions:

- whether the learned trajectories capture biologically meaningful disease progression,
- whether the continuous states reconstructed between sparse observations are predictive,
- and whether the proposed modeling choices are responsible for these properties.

Across these experiments, CohortFM consistently produced patient-specific trajectories that revealed structure not visible when the data were analyzed only through the original clinical visits. One of the clearest findings is that calendar time and molecular progression are not interchangeable. Patients sampled at the same nominal visit frequently occupied different molecular states, while samples collected at different visits could correspond to similar stages of progression. In the COVID-19 cohort, nearly half of the observed samples mapped away from the molecular position most commonly associated with their clinical visit, showing that the model captures differences in how rapidly and in what manner individual patients progress.

This reorganization of the same observed samples substantially increased biological resolution. Compared with the original visit-level representation, CohortFM recovered many more significant biological processes and exposed transient and non-monotonic programs that were otherwise hidden between measurements. In particular, cell-cycle, mitochondrial, translation, and immune-related processes emerged at intermediate progression states, and several shared biological programs showed different timing across trajectories. Thus, the continuous representation does not simply create a smoother version of the original data; it can change the biological interpretation by revealing when a process becomes active, disappears, or re-emerges along a particular disease course.

The cohort-level trajectories also captured meaningful heterogeneity between patients. Even after accounting for a patient’s position along progression, trajectory identity contributed additional information about held-out gene-expression profiles across all four diseases, indicating that patients at similar stages can still follow molecularly distinct courses. Consistent with this observation, CohortFM produced a larger proportion of biologically distinguishable trajectories across the evaluated diseases while retaining all eligible patients in the trajectory representation.

The learned trajectories were also supported from a machine-learning perspective. When asked to reconstruct an unobserved visit between two known observations, CohortFM remained competitive with strong interpolation baselines, while the advantage became clearer in the more difficult task of predicting a future molecular state that was not available to the model. Across the four diseases, CohortFM achieved the strongest patient-level prediction in three cohorts and the strongest cohort-level distributional reconstruction in three. Additional experiments that systematically removed different parts of the training design showed that the model’s performance does not come from a single component alone. Instead, the strongest and most consistent results were obtained when the model combined continuous trajectory learning with additional constraints from observed patient states. This supports the overall design of CohortFM as a model that learns smooth longitudinal dynamics while remaining grounded in real measurements.

Our results show that CohortFM is not simply an interpolation method for sparse observations. It learns a patient-aware representation of molecular progression that separates biological change from sampling time, captures heterogeneous disease courses, reveals transient biological programs that are missed by coarse visit-based analysis, and remains useful for reconstructing and predicting unobserved molecular states.

## 6 Model Design and Trajectory Construction

### 6.1 Variational Autoencoder for Representation Learning

To obtain compact, non-linear representations of the transcriptomic profiles, we use a variational autoencoder (VAE). Each input sample is a 2,000-dimensional gene-expression vector obtained after preprocessing. The purpose of this stage is to compress the high-dimensional molecular profile into a lower-dimensional latent state that preserves the dominant structure of the expression data. The encoder consists of fully connected layers followed by layer normalization, LeakyReLU activation, and dropout. Its final layer produces a latent mean (***µ***) and log-variance (log ***σ***^2^). During training, latent vectors are sampled using the reparameterization trick (**z** = ***µ*** + ***σ*** ⊙ ***ε*** where ***ε*** ∼ *N* (**0, I**)).

The decoder mirrors the encoder and reconstructs the standardized gene-expression profile from the latent representation. The principal architecture uses two hidden layers of 512 and 256 units, with a dropout rate of 0.1. During architecture selection, smaller alternatives were also evaluated where appropriate. The training objective combines expression reconstruction, optional auxiliary clinical supervision, and latent-space regularization, as shown in Equation 1:

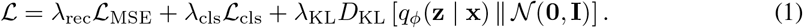

The reconstruction term (*L*_MSE_) minimizes the error between the input and reconstructed expression profiles. An optional auxiliary classification loss (*L*_cls_) can encourage the latent representation to retain clinically relevant information that reconstruction alone may overlook. This component is dataset-dependent, as some datasets provide additional clinical metadata that may enrich representation learning; for example, the sepsis dataset includes in-hospital mortality information. We include this term to keep the framework general and adaptable across datasets, but in all experiments reported here we set *ω*_cls_ = 0, so the latent representations were learned without clinical supervision.

The loss function has a third component, the KL-divergence term (*D*_KL_), to regularize the approximate posterior toward a standard normal distribution. This was designed to produce a smoother, more structured latent space that generalizes better to unseen samples. The auxiliary classification and KL terms were introduced gradually during the first 50 training epochs so that early optimization was driven primarily by reconstruction before the regularization terms reached their full weights.

The VAE is optimized using AdamW [Loshchilov and Hutter, 2019], with an initial learning rate of 10^−3^ and weight decay of 10^−4^. A `ReduceLROnPlateau` scheduler [Al-Kababji et al., 2022] reduces the learning rate when validation performance plateaus, and early stopping is applied with a patience of 40 epochs. Candidate architectures, latent dimensions, and loss weights were selected according to validation reconstruction performance. Batch size was 64 for COVID-19 and 32 for the other three cohorts. The selected latent dimensionalities were 64 for COVID-19 and influenza and 32 for sepsis and tuberculosis. The standard 512–256 hidden-layer architecture was selected for COVID-19, sepsis, and tuberculosis, whereas the influenza architecture search additionally considered a smaller encoder and ultimately selected a single 256-unit hidden layer. For downstream trajectory modeling, we use the posterior mean (**z**_*i*_ = ***µ***_*ϕ*_(**x**_*i*_)) as the deterministic latent representation.

### 6.2 Patient Trajectory Modeling via Flow Matching

Given the low-dimensional representations from the VAE, we model longitudinal disease progression with a flow-matching model we call CohortFM. Its several design choices are tailored to sparse longitudinal patient trajectories, with the objective of learning a continuous, patient-specific trajectory between observed transcriptomic states. For patient *p* at visit *k*, **z**_*p,k*_ denotes the latent representation of the corresponding transcriptomic profile. Visits are ordered chronologically for model training, but calendar time alone does not necessarily reflect the rate of molecular progression. Two patients may undergo substantially different transcriptomic changes over the same period of time; for example, Day 2 for patient *p* may not correspond to the same disease stage as Day 2 for patient *q*. We therefore introduce a normalized pseudotime coordinate *s*_*p,k*_∈ [0, 1] computed from the cumulative distance traveled in the learned latent space.

For consecutive observations, we first compute the distance, as in Equation 2:

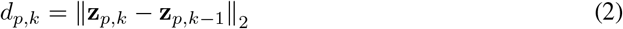

and define the pseudotime of visit *k* as the normalized cumulative distance:

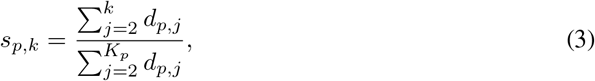

where *K*_*p*_ is the number of observed visits for patient *p*, so that *s*_*p*,1_ = 0 and *s*_*p,Kp*_ = 1. By construction the coordinate is monotonically increasing on [0, 1]. If a trajectory shows almost no change in latent space, the positions are assigned uniformly instead. Calendar time is still important for preserving the temporal structure of the data, so we retain it separately and provide it to the model alongside pseudotime. Pseudotime in our setup complements rather than replaces the actual temporal information.

Training examples are constructed independently within each patient trajectory. For an ordered sequence [**z**_*p*,1_,. .., **z**_*p,Kp*_], we distinguish *short-hop* from *long-hop* transitions to provide complementary supervision at different temporal scales. A short hop is a transition between two immediately consecutive observed visits, and a long hop connects non-adjacent visits. For example, if a patient has three consecutive sampling points, the transition from Visit 1 to Visit 2 is a short hop, while the transition from Visit 1 to Visit 3 is a long hop. Such a patient therefore contributes to training by two short hops (Visit 1→Visit 2 and Visit 2→Visit 3) and one long hop (Visit 1→Visit 3).

Short hops, collected in the set *ε*_short_, give direct supervision for local transitions; long hops, collected in *ε*_long_, expose the model to longer-range progression. Because short hops are directly observed local transitions, they are assigned unit weight, whereas long hops are down-weighted (*ω*_long_ *<* 1). Long hops remain informative for global trajectory structure but span larger intervals and so they could carry greater uncertainty in the implied path. The long-hop weight is a hyper-parameter selected on the validation cohort to account for this. In this way, long-range transitions improve global structure without overwhelming the more reliable local supervision.

For a transition from Visit *i* to *j*, we retain both the calendar-time displacement (Δ*t*_*ij*_ = *t*_*p,j*_ *t*_*p,i*_) and the pseudotime displacement (Δ*s*_*ij*_ = *s*_*p,j*_ *s*_*p,i*_). Later paragraphs describe how the model uses them. Because transition durations could vary, we scale Δ*t* by the median training-set transition duration, resulting in 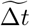.

#### 6.2.1 Flow-Matching Model Training

The training strategy has three components: *i*) a flow-matching loss, *ii*) a path-consistency loss, and *iii*) a velocity-anchor loss. Together they train a flow-matching model that captures patient-specific longitudinal structure, rather than acting as a simple generative interpolator. For each observed transition (*i, j*), we denote its endpoints **z**_0_ = **z**_*p,i*_ and **z**_1_ = **z**_*p,j*_. During training we sample a local bridge coordinate τ ∼ *U* (0, 1), where *τ* = 0 is the source visit and *τ* = 1 the target. *τ* is local to a single transition, in contrast to the global patient pseudotime *s*. The intermediate latent state is the linear interpolation **z**_*τ*_ = (1 − *τ*)**z**_0_ + *τ* **z**_1_, with pseudotime *s*_*τ*_ = (1 − *τ*)*s*_0_ + *τs*_1_. The model is trained to predict the direction from source to target, **z**_1_ − **z**_0_, so that it learns from positions throughout each transition rather than from the endpoints alone. We sampled two such positions per transition per training epoch.

CohortFM learns a neural vector field **v**_*θ*_, as explained in Equation 4:

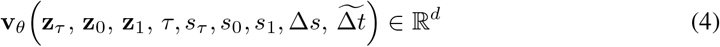

Rather than conditioning only on the current state and time, the model is conditioned on both endpoints. The full input comprises the current state **z**_*τ*_ ; source and target states **z**_0_, **z**_1_; the local position *τ* ; the pseudotimes *s*_*τ*_, *s*_0_, *s*_1_ displacement Δ*s*: and and the normalized calendar-time displacement 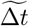.

Because *τ* represents position within a transition, using it as a single scalar may be too limited. We therefore augment *τ* with sinusoidal features at multiple frequencies. Specifically, for *ω* ∈*{*1, 2, 4, 8 *}*, we compute sin(2*πωτ*) and cos(2*πωτ*) and concatenate these eight values with *τ* and the remaining model inputs. Lower frequencies capture coarse progression across the interval, while higher frequencies represent finer variations in the learned vector field. Taken together, these features give an input dimensionality of 3*d* + 14 for a *d*-dimensional latent space.^6^

The network uses an initial fully connected projection layer, followed by a series of residual MLP blocks. Each block applies layer normalization, a hidden expansion, SiLU activation [Elfwing et al., 2018], dropout, and projection back to the model dimension, with a residual connection. A final linear layer outputs the predicted latent velocity.

##### Flow-matching loss

The primary objective minimizes the discrepancy between predicted and reference velocities:

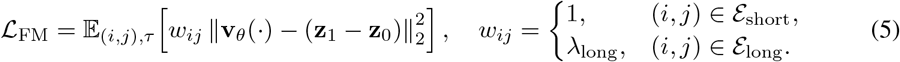

This learns a cohort-level vector field from all patient transitions while preserving the distinction between strongly supervised local transitions and weaker long-range ones.

#### Path-consistency loss

Flow matching can favor nearly straight paths between endpoints, whereas real patient trajectories may be nonlinear. To encourage more realistic paths, we use observed intermediate visits as additional supervision. Consider a long transition from **z**_0_ to **z**_1_ with an observed intermediate state **z**_*h*_ at pseudotime *s*_*h*_. Its relative position within the transition is defined in Equation 6:

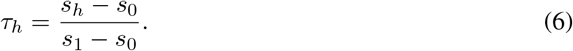

Starting from **z**_0_, we integrate the learned vector field up to *τ*_*h*_. Let *m* denote the numerical integration step, with **z**^(*m*)^ representing the latent state at step *m*. The update is given in Equation 7:

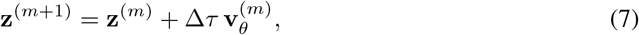

where Δ*τ* is the integration step size and **v**_*θ*_ is defined in Equation 4. Let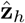 denote the state reached at *τ*_*h*_. We then encourage this predicted state to match the observed intermediate state using the path-consistency loss in Equation 8:

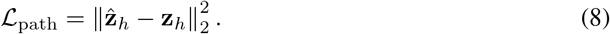

For example, for three consecutive visits, the model considers the long transition from Visit 1 to Visit 3, starts from Visit 1, and integrates the learned vector field up to the relative position corresponding to the observed intermediate Visit 2. The resulting latent state is then penalized if it differs from the true Visit 2 representation. This encourages the learned trajectory between Visit 1 and Visit 3 to pass through the observed intermediate state rather than defaulting to a simple linear interpolation.

##### Velocity-anchor loss

We further use observed intermediate states as local velocity anchors. For an intermediate state **z**_*h*_ in a long transition, we estimate incoming and outgoing local velocities, as shown in Equation 9:

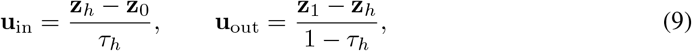

and define the central velocity target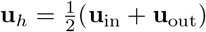. The vector field at the observed intermediate state is encouraged to match this local tangent:

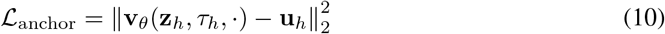

This adds local directional supervision alongside the integrated path-consistency constraint.

##### Final loss

The training objective combines the three components, as in Equation 11:

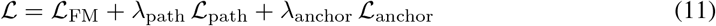

where *τ*_path_ and *τ*_anchor_ balance the path-consistency and anchor terms against the primary flow-matching loss and are tuned over a validation set during training. To prevent leakage between repeated observations of the same individual, the development split was at the *patient* level: patients were randomly divided with 20% assigned to validation, and all of a patient’s visits and transitions kept in the same partition. We ran a compact hyper-parameter search over the width and number of residual blocks of the network described above as well as other hyper-parameters. Models were optimized with AdamW at learning rate 10^−3^, weight decay 10^−4^, batch size 64, dropout 0.08, and gradient clipping at norm 1.0.

#### 6.2.2 Constructing Patient Trajectories

After training, CohortFM reconstructs each patient transition by following the learned vector field between consecutive observed states, producing a continuous latent trajectory between visits. Since the vector field is conditioned on both endpoints, pseudotime, and calendar duration, the resulting path remains specific to each patient interval. For trajectory augmentation, we generate one latent midpoint between adjacent visits by integrating to *τ* = 0.5, rather than taking a simple linear average.

Beyond generating intermediate states, the pseudotime coordinate provides a common scale for comparing patients with different temporal dynamics. Each trajectory is evaluated on the normalized grid *S* = *{*0, 0.2, 0.4, 0.6, 0.8, 1.0*}*. For each position *s*^∗^∈ *S*, an observed latent state is used directly when available; otherwise, the learned flow is integrated between the neighboring observed states to generate the state at *s*^∗^. This produces trajectories with the same number of normalized states while preserving each patient’s observed measurements and learned trajectory shape. Examples of these trajectories from our COVID-19 dataset are shown in Figure 2, providing a more tangible illustration of the concept.

**Figure 2.**
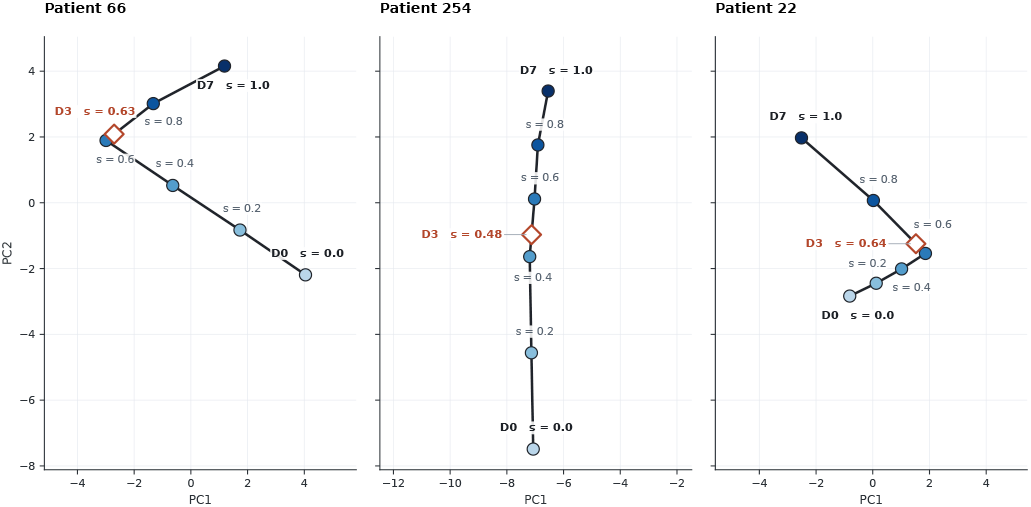
Reconstructed trajectories for Patients 66, 254, and 22. Each patient progresses through the same normalized molecular milestones on a shared coordinate system and scale, allowing their disease-progression patterns to be compared directly. For visualization, the original *d*-dimensional trajectories are projected into two dimensions.

### 6.3 Mixture Modeling of Cohort-Level Trajectories

To move from patient-specific trajectories to a cohort-level representation, we need a small set of trajectories that capture the dominant longitudinal patterns across patients. We therefore introduce a mixture model that operates downstream of CohortFM on the normalized patient trajectories, without modifying the learned patient-level flow model.

Let _*p*_ denote the normalized latent trajectory of patient *p* on the shared grid *S*, as illustrated in Figure 2. The model learns *T* cohort-level trajectories [**T**_1_,. .., **T**_*T*_], where each **T**_*t*_ represents a characteristic progression pattern in the cohort.^7^ Rather than assigning each patient to only one of these trajectories, the model allows a soft association with several of them. For each patient trajectory *P*_*p*_, the encoder produces one score for each cohort-level trajectory, and a softmax converts these scores into probabilities. The patient trajectory is then reconstructed as a probability-weighted combination of the learned cohort-level trajectories, as shown in Equation 12:

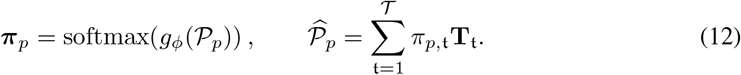

where *g*_*ϕ*_ is the mixture encoder, parameterized by *ϕ*, and *π*_*p,t*_ is the probability that patient *p* is associated with cohort-level trajectory **T**_*t*_. The probabilities sum to one across the *T* trajectories, and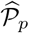is the reconstructed patient trajectory obtained from their weighted combination. A patient is therefore not restricted to a single cohort-level trajectory. Instead, the model can represent each patient as a mixture of several progression patterns. Once training is complete, each patient is assigned to the cohort-level trajectory with the highest probability.

The mixture encoder uses two fully connected transformations with ReLU activations and dropout, followed by a -dimensional output layer. The cohort-level trajectories **T**_*t*_ are learnable parameters, initialized by clustering the flattened patient trajectories to provide a stable starting point for joint *T*optimization. The primary reconstruction objective is defined in Equation 13:

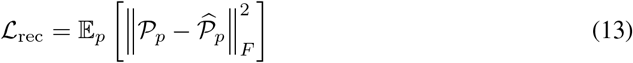

which encourages the learned cohort-level trajectories and mixture probabilities to jointly reconstruct the patient trajectories. Figure 3 visualizes the output of the mixture model.

**Figure 3.**
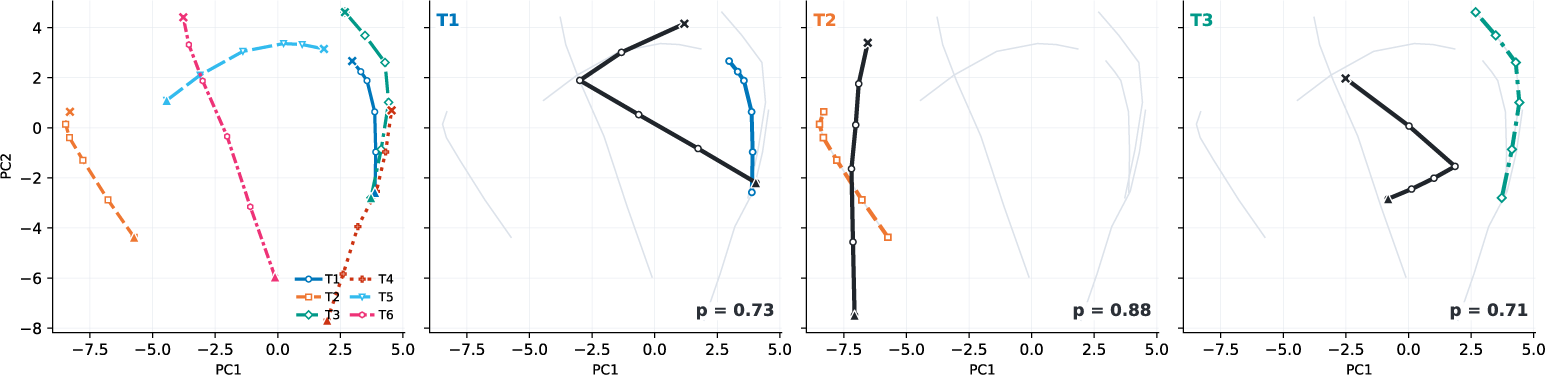
Learned cohort-level trajectories and example patient assignments. The first panel shows a real example from our COVID-19 dataset, where the full cohort is distilled into six dominant cohort-level progression trajectories. The remaining panels show three example patient-specific trajectories mapped to the cohort-level trajectory with which they are most strongly associated. The solid black trajectories are the same patient trajectories shown in Figure 2.

Once training is complete, the model yields *T* cohort-level trajectories that compactly summarize the major patient trends, progression patterns, and longitudinal behaviors observed across the cohort. These trajectories form the basis for subsequent biological analysis and evaluation. At this stage, the modeling pipeline is complete, and the focus shifts to interpreting and evaluating the learned cohort-level structure.

As with any mixture model, reconstruction accuracy alone does not guarantee that the learned trajectories are useful. Without additional constraints, some trajectories may become redundant, be used by very few patients, or become unnecessarily irregular. We therefore add four regularization terms that encourage smooth trajectories, balanced use of the cohort-level trajectories, and diversity across them. These terms help stabilize mixture-model training. Full definitions of these complementary loss components are provided in Appendix B.

### 6.4 Mapping Patients to Cohort-Level Trajectory States

Once the cohort-level trajectories have been learned, the next step is to connect individual patients and their observed samples back to these trajectories so that the resulting progression patterns can be analyzed biologically. Each patient is first assigned to the cohort-level trajectory with the highest mixture-model probability. We then consider each observed sample from that patient and determine where it lies along the assigned trajectory. Specifically, the sample is compared with the normalized positions along the trajectory and mapped to the position with the most similar latent molecular state.

This process groups observed samples according to both their cohort-level trajectory and their position along that trajectory. We refer to the samples grouped around the same trajectory position as a *patient cloud*. Each cloud therefore represents a set of observed patient samples occupying a similar molecular state within a particular progression pattern. Figure 4 visualizes this concept.

**Figure 4.**
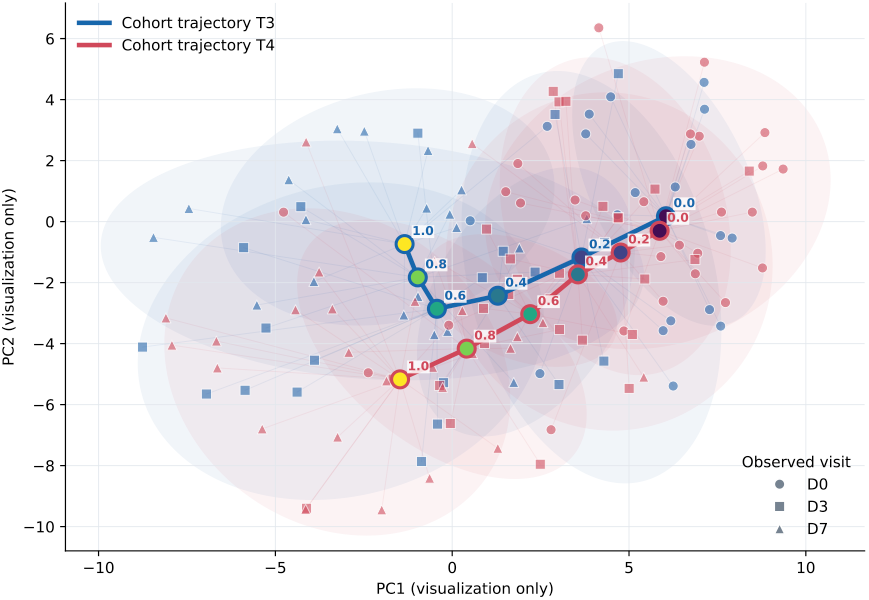
Illustration of patient-cloud construction for two cohort-level trajectories from the COVID-19 dataset. Observed patient samples are mapped to their closest normalized position along the assigned trajectory, forming local clouds around different progression states. Marker shapes indicate the original clinical visits. The two-dimensional projection is shown only for visualization; sample-to-trajectory mapping is performed in the full latent space.

Importantly, this mapping is performed entirely in the learned latent molecular space and does not use calendar visit labels. Samples collected at the same clinical visit can therefore map to different progression states, while samples collected at different visits can occupy similar molecular states. These patient clouds connect the learned trajectory representation back to the original biological measurements. The corresponding molecular profiles can then be used for downstream analyses such as characterization of biological programs along each cohort-level trajectory.

## 7 Detailed Evaluation Results

This section provides a comprehensive evaluation of CohortFM using the COVID-19 dataset as our primary case study. We use this cohort to examine the learned trajectories in detail, both mathematically and biologically, and to characterize the strengths and limitations of the framework. CohortFM is also evaluated on three additional longitudinal datasets, but repeating the full set of analyses for every dataset would be unnecessarily extensive. We therefore use the COVID-19 dataset for the detailed analysis and present a more compact, standardized evaluation of the remaining datasets in Section 8.

### 7.1 Do the Learned Trajectories Provide a Meaningful Representation of the Cohort?

We first examine whether the learned cohort-level trajectories provide a meaningful representation of the COVID-19 cohort and are sufficiently populated for downstream biological analysis. CohortFM identifies six progression trajectories, denoted **T**_1_–**T**_6_, with 20, 10, 20, 28, 13, and 32 patients assigned to them, respectively, for a total of 123 patients. During training, CohortFM can leverage nearly all available longitudinal observations. Once the cohort-level trajectories are learned, each patient is associated with the trajectory for which they have the highest mixture-model assignment probability. The patient counts above therefore reflect the dominant trajectory assignment within the analysis subset rather than an imposed partition of equal size.

Although the full COVID-19 cohort contains 306 positive patients, we use the 123 patients with complete D0, D3, and D7 measurements for biological characterization of the learned trajectories. This restriction is applied *only* to the downstream analysis and *does not* limit the observations available during trajectory learning. Using complete longitudinal profiles provides a cleaner and more comparable basis for biological interpretation, which is particularly important in studies where cohort sizes are already limited. In principle, patients with incomplete follow-up (e.g., only D0 and D3) can also be mapped to the learned cohort-level trajectories.

The mixture model is expected to form meaningful groups with sufficient support for downstream analysis. Extremely small trajectories, such as those containing only one or a few patients, or a single trajectory dominating most of the cohort, would need further investigation. Here, all six trajectories are supported by a meaningful number of patients, with none collapsing to a few observations or dominating the cohort. This provides a sufficiently populated basis for the trajectory-level biological analyses that follow.

Having established a set of well-supported cohort-level trajectories, we next examine how the original calendar visits are distributed along the normalized molecular-progression axis. Figure 5a shows the pooled composition of D0, D3, and D7 samples across the normalized positions *s*∈ *{*0, 0.2, 0.4, 0.6, 0.8, 1.0*}*. If calendar time and molecular progression were interchangeable, samples from the same visit would be expected to concentrate at similar positions. Instead, the observed visits are distributed across multiple molecular progression states. For example, in **T**_1_, all D0 observations occur at *s* = 0, a pattern also observed in **T**_3_ and **T**_4_. In **T**_5_, however, approximately 8% of the samples assigned to the initial molecular position (*s* = 0) are D3 observations rather than D0. Even around 8% of the D7 samples are also in the same position. These differences illustrate that the nominal visit label alone does not uniquely determine where a patient lies along the molecular progression axis. The complete per-trajectory breakdown is provided in Figure 12 in Appendix C.

**Figure 5.**
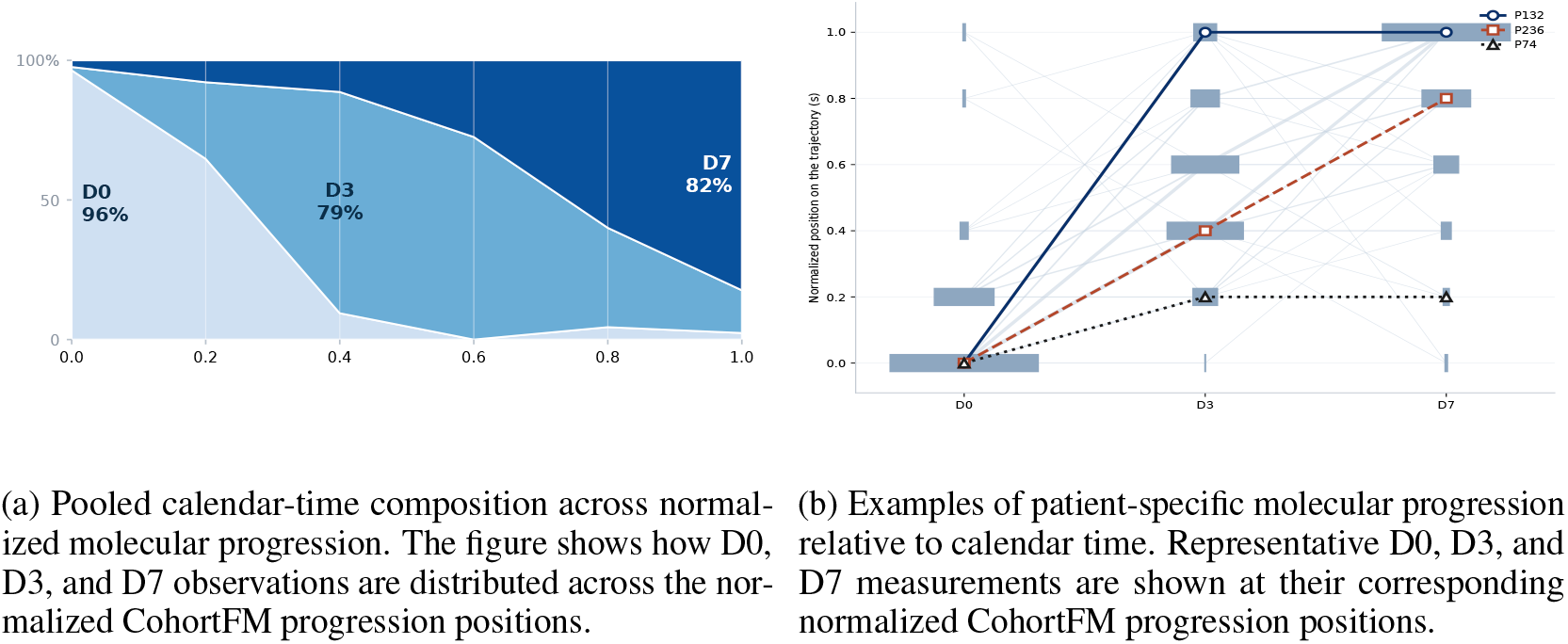
Calendar time versus normalized molecular progression. (a) Samples collected at the same nominal clinical visit can occupy different molecular progression states. (b) Individual examples further demonstrate that patients sampled at the same calendar time can exhibit substantially different amounts of molecular progression.

This distinction becomes clearer when individual patients are examined. Figure 5b shows representative examples of how the three observed visits from individual patients map onto the normalized progression coordinate. Patient 74 (triangle) begins at *s* ≈ 0 at D0 and progresses to approximately *s* ≈0.2 by D3, but remains near the same molecular position at D7. Additional elapsed time therefore need not correspond to additional molecular progression. Patient 132 (circle) follows a markedly different course. The D0 sample lies near the beginning of the trajectory, whereas both the D3 and D7 samples occur close to the end of the normalized progression axis. Patient 236 (square) exhibits a more gradual pattern, with D0, D3, and D7 mapping approximately to *s* = 0, 0.4, and 0.8, respectively.

These examples capture one of the central motivations for CohortFM. Clinical sampling schedules provide a common temporal reference, but they do not imply that different patients have progressed by the same molecular amount. By representing calendar time and molecular progression separately, CohortFM provides both a trajectory identity and a relative molecular position for each patient. This enables patients with different rates and patterns of progression to be compared within a common coordinate system while preserving their individual longitudinal behavior.

### 7.2 Biological Programs Along Cohort-Level Trajectories

A central goal of CohortFM is to determine whether the learned cohort-level trajectories capture meaningful biological progression rather than only geometric structure in latent space. If they do, patient clouds at different positions along a trajectory should correspond to coherent molecular states, and those states should vary systematically along normalized progression. We therefore mapped each patient cloud back to gene-expression space and characterized the biological processes represented by the real patient samples within it.

For each patient cloud, gene expression was compared with the control group, followed by GO Biological Process enrichment of significantly altered genes.^8^ Across all the patient clouds, this analysis identified 1,277 significant cloud–GO associations spanning 396 unique GO terms. Because many GO terms describe closely related processes, we consolidated them into seven broader biological programs: antiviral defense and interferon, innate and antimicrobial response, adaptive immune signaling, cytokine and inflammatory signaling, cell cycle and mitosis, mitochondrial energetics, and translation and ribosome. Overall, 74.9% of the significant GO associations could be assigned to one of these programs.

Figure 6 shows how these biological programs are distributed along the six cohort-level trajectories. Each panel follows one trajectory from *s* = 0 to *s* = 1, allowing the biological signal to be examined not only across trajectories but also across different stages of molecular progression. The figure summarizes enrichment among genes elevated relative to the control group, with stronger colors indicating stronger GO enrichment within the corresponding patient cloud.

**Figure 6.**
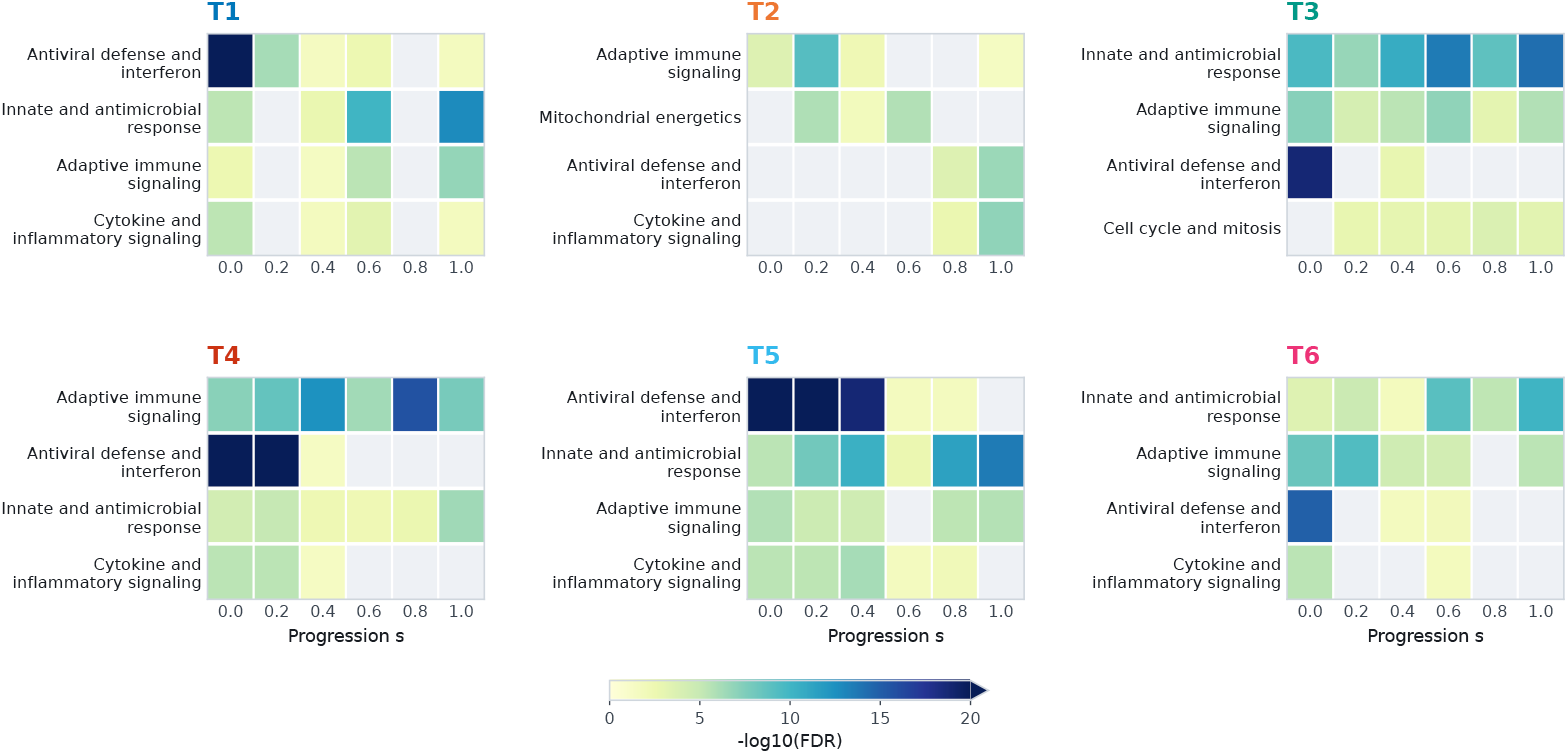
Biological programs along the learned cohort-level trajectories. Each panel represents one cohort-level trajectory, with normalized molecular progression *s* from 0 to 1. Rows show the dominant biological programs identified from the patient clouds along that trajectory. Color represents the log_10_(FDR) of the strongest significant GO Biological Process term associated with genes elevated relative to the control group; blank cells indicate that no term from the corresponding program reached the significance threshold.

The learned trajectories are therefore biologically interpretable: several immune-related programs recur across the cohort, while their strength and location along progression vary between trajectories.

The trajectory representation therefore provides more than a grouping of molecularly similar patients. It organizes real biological signals according to both the type of progression followed by a patient and the patient’s position along that progression.

### 7.3 Distinct Biological Profiles Across Cohort Trajectories

As shown in the previous section, the patient clouds capture coherent biological signals. We next investigate how these signals differ across the six cohort-level trajectories. Table 2 summarizes the dominant biological programs associated with each trajectory, the progression positions at which they are detected, and representative genes that most strongly distinguish each trajectory from the others.

**Table 1.** Summary of the longitudinal transcriptomic datasets used in our experiments.

| Dataset | Accession | Assay / Tissue | Patients | Longitudinal Sampling |
| --- | --- | --- | --- | --- |
| COVID-19 | GSE212041 | Bulk RNA-seq / Neutrophils | 306 COVID-19<sup>+</sup> | D0, D3, D7 |
| Sepsis<sup>5</sup> | GSE54514 | Microarray / Whole blood | 36 | Daily, up to 5 days |
| Severe Influenza | GSE111368 | Microarray / Whole blood | 109 | T1, T2 (~2 days), T3 (recovery) |
| Tuberculosis | GSE89403 | Bulk RNA-seq / Whole blood | 98 | Week 0, 1, 4, 24 |

**Table 2.** Biological profiles of the six cohort-level trajectories. For each trajectory, the table reports its dominant biological programs, the normalized progression positions at which those programs are significantly detected, and representative genes. Gene values in parentheses are the log_2_ fold-change contrasts used to identify genes that distinguish each trajectory from the others.

| Dominant biological programs | Progression positions | Representative genes |
| --- | --- | --- |
| **T1:** Antiviral defense and interferon; Innate and antimicrobial response; Adaptive immune signaling | Antiviral: $s = 0.0-0.6, 1.0$; Innate: $s = 0.0, 0.4-0.6, 1.0$; Adaptive: $s = 0.0, 0.4-0.6, 1.0$ | *PI3* (+1.14), *CXCL2* (+0.76), *CXCL8* (+0.69), *IFITM3* (+0.60) |
| **T2:** Adaptive immune signaling; Mitochondrial energetics; Antiviral defense and interferon | Adaptive: $s = 0.0-0.4, 1.0$; Mitochondrial: $s = 0.2-0.6$; Antiviral: $s = 0.8-1.0$ | *CD8A* (+2.15), *LCK* (+1.95), *RSAD2* (+1.89), *IFIT1* (+1.84) |
| **T3:** Innate and antimicrobial response; Adaptive immune signaling; Antiviral defense and interferon | Innate: $s = 0.0-1.0$; Adaptive: $s = 0.0-1.0$; Antiviral: $s = 0.0, 0.4$ | *DEFA1* (+3.33), *CTSG* (+2.65), *PRTN3* (+2.59), *DEFA4* (+2.58) |
| **T4:** Adaptive immune signaling; Antiviral defense and interferon; Innate and antimicrobial response | Adaptive: $s = 0.0-1.0$; Antiviral: $s = 0.0-0.4$; Innate: $s = 0.0-1.0$ | *IGHG1* (+1.17), *IGHG3* (+1.13), *IGHA1* (+1.13), *IGLC3* (+1.09) |
| **T5:** Antiviral defense and interferon; Innate and antimicrobial response; Adaptive immune signaling | Antiviral: $s = 0.0-0.8$; Innate: $s = 0.0-1.0$; Adaptive: $s = 0.0-0.4, 0.8-1.0$ | *RSAD2* (+1.75), *IFIT3* (+1.68), *IFIT1* (+1.64), *ISG15* (+1.56) |
| **T6:** Innate and antimicrobial response; Adaptive immune signaling; Antiviral defense and interferon | Innate: $s = 0.0-1.0$; Adaptive: $s = 0.0-0.6, 1.0$; Antiviral: $s = 0.0, 0.4-0.6$ | *IGHG1* (+1.41), *IGKV3-20* (+1.23), *CXCL8* (+1.16), *IGLC3* (+1.12) |

The six trajectories capture different patterns of shared immune biology. **T**_3_ is characterized by persistent innate and adaptive immune programs across the full progression axis and has the strongest aggregate innate and antimicrobial signal, with *DEFA1, CTSG, PRTN3*, and *DEFA4* among its most distinguishing genes. **T**_4_ also maintains innate and adaptive immune signals throughout progression, but has the strongest aggregate adaptive immune signal and is distinguished by immunoglobulin-related genes (*IGHG1, IGHG3, IGHA1*, and *IGLC3*). **T**_5_, in contrast, has the strongest antiviral and interferon signal, extending from *s* = 0 through *s* = 0.8, with *RSAD2, IFIT3, IFIT1*, and *ISG15* as representative markers.

**T**_2_ shows a different temporal organization. Adaptive immune signaling is concentrated primarily in the early part of the trajectory, mitochondrial energetics appears at intermediate positions (*s* = 0.2– 0.6), and antiviral signaling appears later at *s* = 0.8–1.0. The corresponding GO processes include T Cell Receptor Signaling Pathway, Mitochondrial Translation, and Negative Regulation Of Viral Process. **T**_1_ and **T**_6_ show broader combinations of antiviral, innate, and adaptive programs, but with different coverage along the progression axis. This further illustrates that similar biological processes can be organized differently across patient progression patterns.

We next asked which biological programs are broadly shared across trajectories and which are more trajectory-specific. Cell cycle and mitosis, innate and antimicrobial response, antiviral defense and interferon, adaptive immune signaling, and cytokine and inflammatory signaling were all shared across multiple trajectories. Their strongest signals nevertheless occurred in different trajectories. Cell-cycle and innate programs were strongest in **T**_3_, antiviral and cytokine/inflammatory programs in **T**_5_, and adaptive immune signaling in **T**_4_. In contrast, mitochondrial energetics and translation/ribosome were confined to **T**_2_ in the above-control enrichment signal, each receiving the maximum trajectory-specificity score of 1.0.

Together, these results show that the six cohort-level trajectories are not redundant versions of the same disease response. Much of the immune biology is shared across the cohort, but CohortFM separates patients according to how strongly these programs are expressed and where they occur along molecular progression. At the same time, the trajectory representation exposes more selective biological signals that are concentrated within particular progression patterns.

### 7.4 STEM-Style Analysis Reveals Distinct Dynamic Gene Programs Across Trajectories

The preceding analyses characterize which biological programs are associated with different cohort-level trajectories and progression states. We further assessed whether these programs also follow distinct *dynamic expression patterns* along normalized molecular progression. To address this, we performed a STEM-style analysis on gene expression across the six normalized positions of each trajectory. For each gene, we rescaled its six-point expression profile to focus on the pattern of change over progression, then matched that shape to predefined temporal patterns, including increasing (inc), decreasing (dec), early-rise (e-rise), early-drop (e-drop), and late-peak (l-peak) or late-dip (l-dip) profiles. Genes were retained only if they matched a temporal pattern above a fixed correlation threshold and showed sufficient expression change. A trajectory–pattern pair was considered significant only if it contained more genes than expected under a permutation null.^9^

Across the six trajectories, 33 of the 60 possible trajectory–pattern combinations were significant. GO Biological Process enrichment of genes within these patterns identified 160 significant terms across 18 dynamic patterns, which were subsequently summarized into 13 broader biological programs. Figure 7 provides an overview of these dynamic programs across the six trajectories.

**Figure 7.**
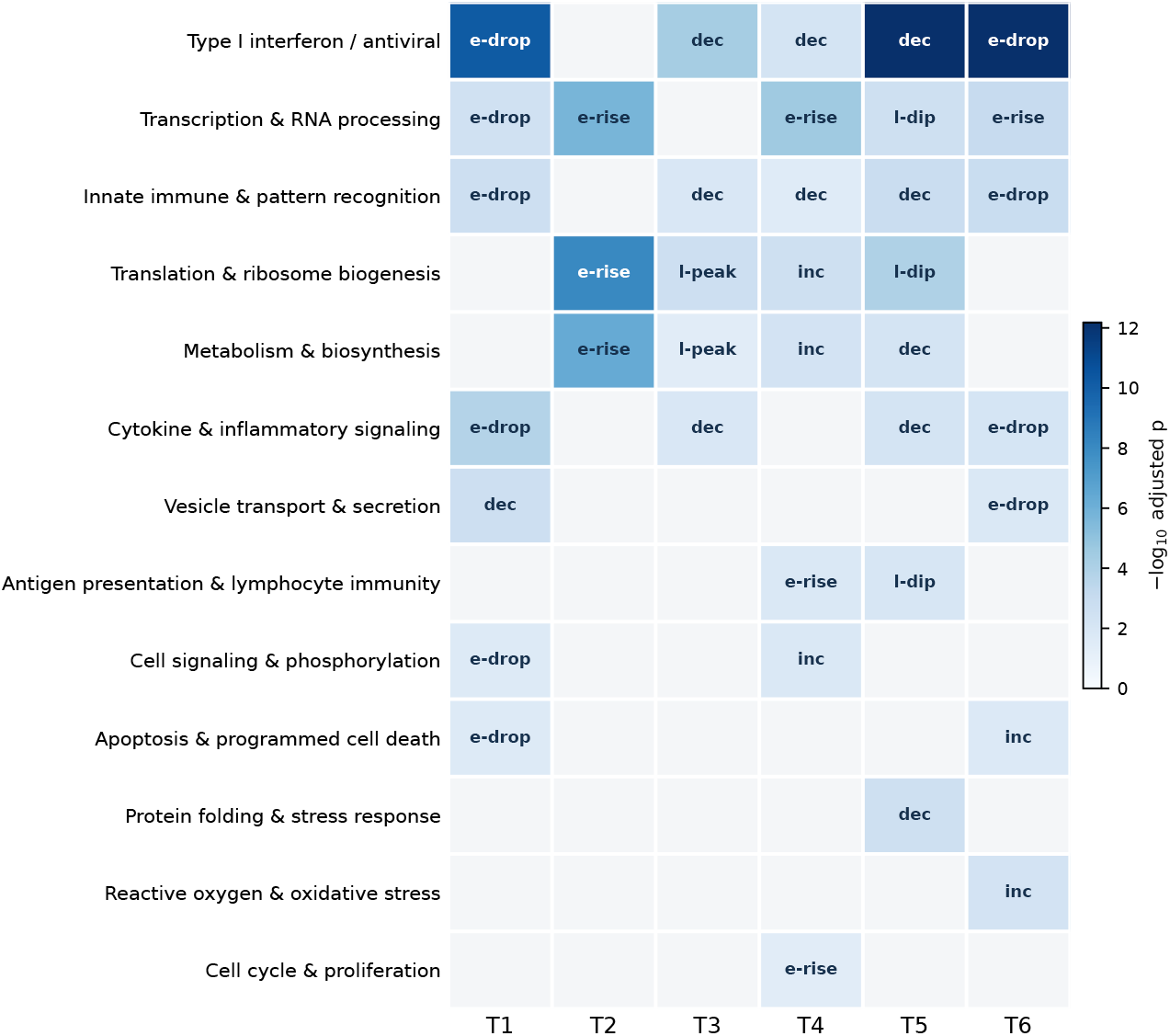
Dynamic biological programs across the six cohort-level trajectories. Rows represent biological programs and columns represent trajectories. Color indicates the significance of the strongest enriched GO Biological Process term, while the label within each cell indicates the corresponding temporal expression pattern. Grey cells indicate trajectory–program combinations without significant support.

The analysis reveals that biological programs can be shared across trajectories while following different progression dynamics. For example, Type I interferon/antiviral and innate immune/pattern-recognition programs were identified in five of the six trajectories. Both followed an early-drop-then-plateau pattern in **T**_1_ and **T**_6_, whereas they showed a decreasing profile in **T**_3_, **T**_4_, and **T**_5_.

Translation and ribosome biogenesis showed even greater variation. It followed an early-rise-then-plateau pattern in **T**_2_, a late peak in **T**_3_, a progressive increase in **T**_4_, and a late dip in **T**_5_. Metabolism and biosynthesis showed a similar contrast, with an early rise in **T**_2_, a late peak in **T**_3_, an increasing profile in **T**_4_, and a decreasing profile in **T**_5_.

The corresponding expression trajectories are shown directly in Figure 8 for a more detailed view. Rather than representing only whether a program is present, these profiles show how the mean standardized expression of its contributing genes changes from *s* = 0 to *s* = 1. The observed curves therefore make explicit that the same biological program can evolve differently depending on the cohort-level progression trajectory. The dynamic analysis also identified more restricted patterns. Antigen presentation and lymphocyte immunity was detected in **T**_4_ and **T**_5_ but followed an early-rise pattern in **T**_4_ and a late-dip pattern in **T**_5_. Several programs were observed in only one trajectory in this analysis. Cell cycle and proliferation followed an early-rise pattern in **T**_4_, protein folding and stress response decreased along **T**_5_, and reactive oxygen and oxidative stress increased along **T**_6_.

**Figure 8.**
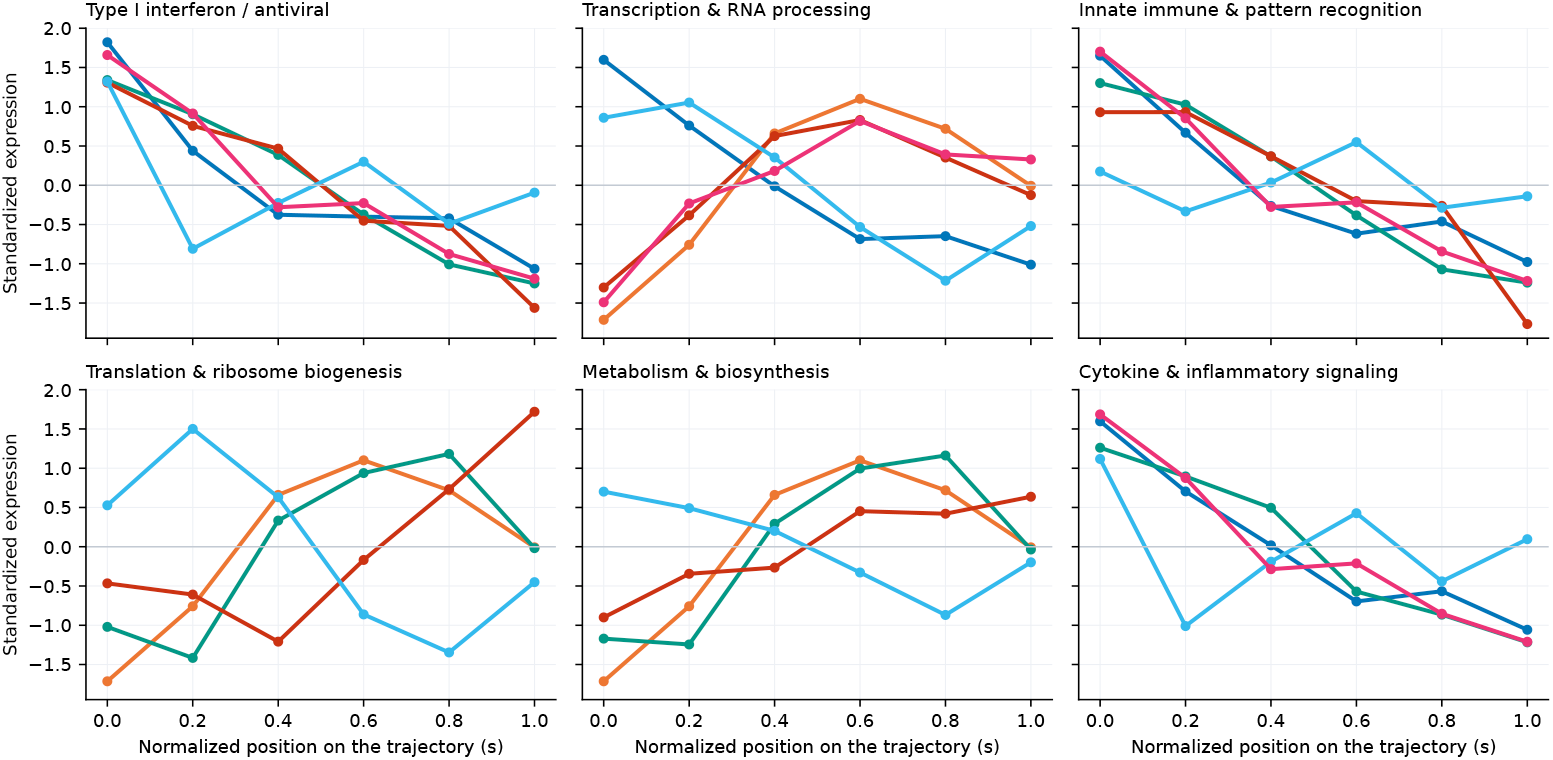
Temporal expression profiles of the leading dynamic biological programs. Curves show the mean standardized expression of genes contributing to each program across normalized molecular progression. A trajectory is shown only when the corresponding program has sufficient gene support.

Results provided in Figures 7 and 8 extend the biological interpretation of CohortFM from *which* molecular programs characterize each trajectory to *how* those programs evolve along progression. This means that even when trajectories involve similar biological processes, those processes can change differently over time, and some programs appear only in specific progression patterns.

### 7.5 Intermediate Molecular States Recover Biological Information

A key motivation for CohortFM is that sparse calendar visits may miss biologically meaningful states that occur between observations. We therefore examined whether CohortFM’s finer molecular-progression scale reveals information beyond the original D0, D3, and D7 measurements. We compared two representations of the same 369 (123 patients ×3 measurements) observed samples: the original three calendar visits and the CohortFM representation consisting of six cohort-level trajectories evaluated at six normalized molecular positions. No synthetic observation was used in this comparison; instead, the learned intermediate states provide a finer coordinate system for organizing the real patient samples. The same analysis thresholds were used in both representations.

Calendar time did not map one-to-one to molecular progression. Overall, 47.7% of samples were assigned away from the most common molecular position of their own visit. Moreover, every molecular position contained samples from more than one calendar visit, and five of the six positions contained samples from all three visits. Thus, patients observed on the same calendar day can occupy different molecular progression states, while samples collected on different days can occupy similar states. This additional resolution substantially enriched the biological analysis (Figure 9).

**Figure 9.**
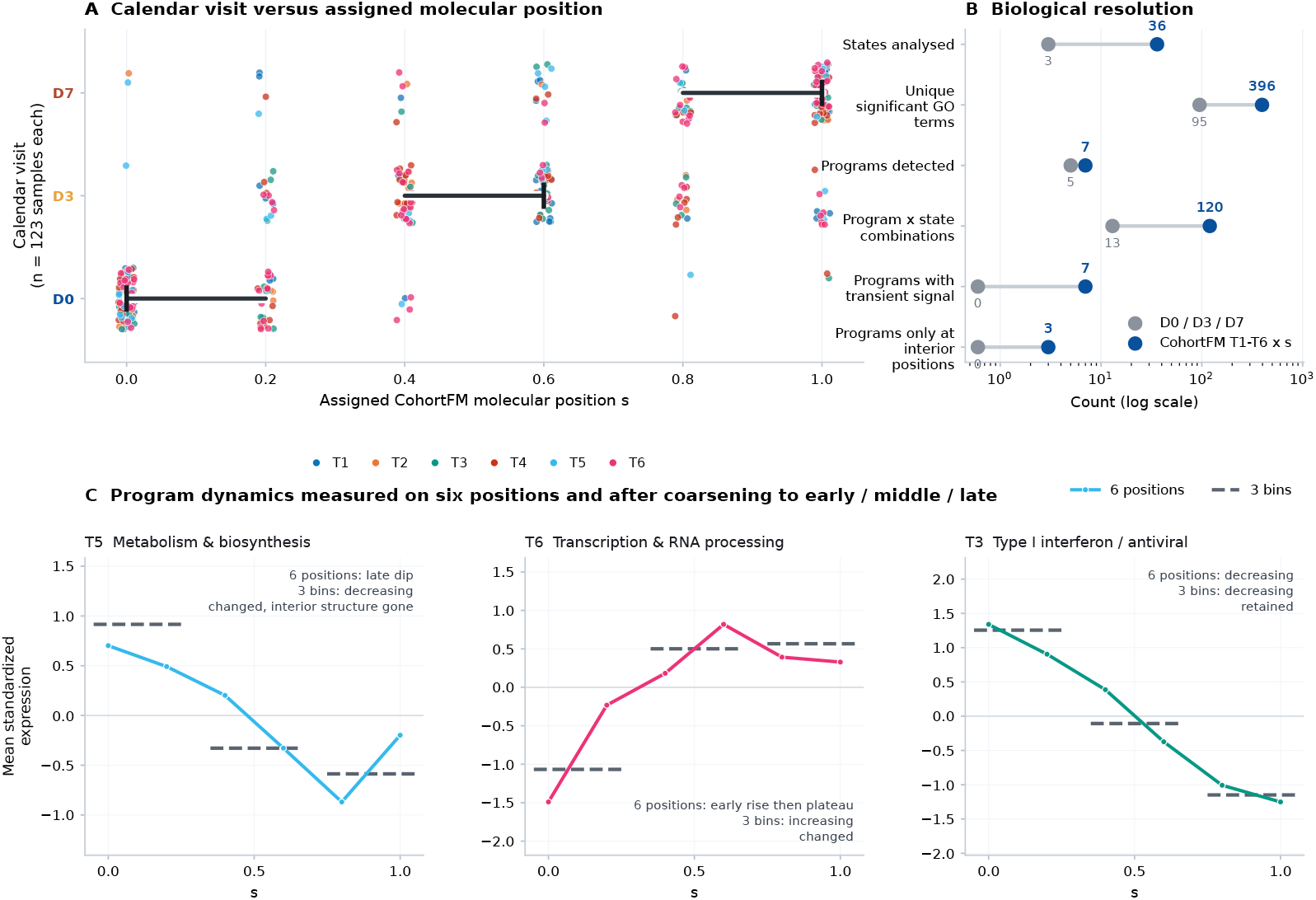
Value of the intermediate molecular states recovered by CohortFM. **A**, Relationship between calendar visit and assigned molecular progression position. Samples collected at the same visit occupy multiple molecular positions, while individual positions can contain samples from different visits. **B**, Biological resolution obtained from the original D0 | D3 | D7 representation and from the CohortFM trajectory–position representation using the same observed samples. **C**, Examples of biological-program dynamics measured at six molecular positions and after deliberately coarsening the same trajectories into early, middle, and late bins. Coarsening can merge or alter temporal patterns that are distinguishable at the full CohortFM resolution.

**Figure 10.**
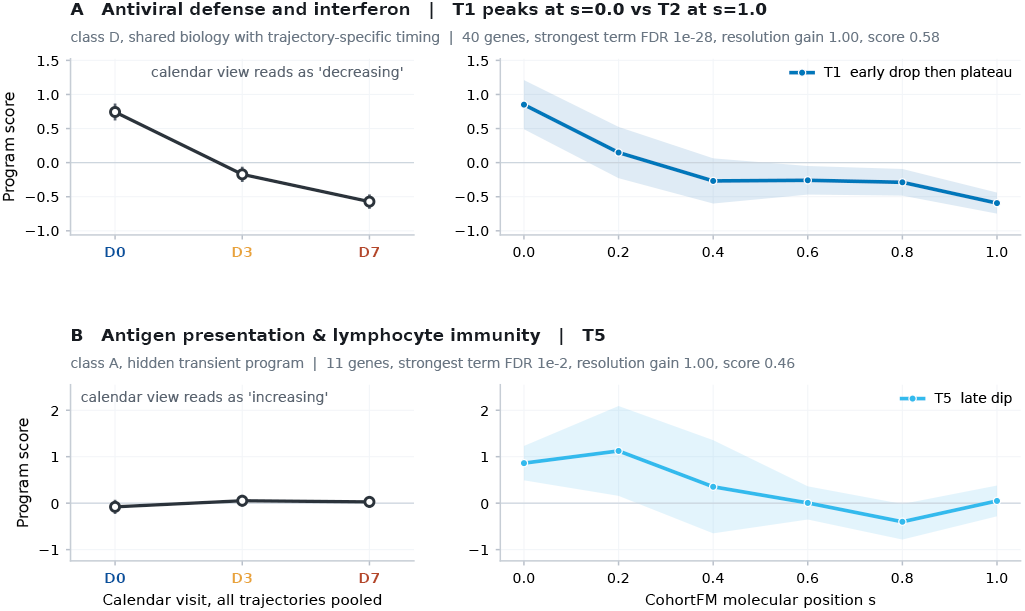
Examples of biological findings enabled by the high-resolution trajectory representation. Left panels show the corresponding calendar-level profiles obtained by grouping the same samples at D0, D3, and D7. Right panels show the biological signal along the CohortFM molecular progression axis. The examples illustrate trajectory-dependent timing and transient biological structure that are obscured when samples are analyzed only at coarse calendar resolution.

The visit-level representation identified 95 unique significant GO terms and five of the seven biological programs, whereas the CohortFM representation identified 396 unique terms and *all seven programs*. More importantly, the number of distinct program–state combinations increased from 13 to 120. The finer representation also revealed transient or non-monotonic behavior in all seven biological programs and identified three programs (cell cycle and mitosis, mitochondrial energetics, and translation and ribosome) that were detectable only at interior progression positions. None of these behaviors was visible in the three-visit analysis.

To isolate the contribution of the intermediate positions themselves, we kept the trajectory assignments fixed and deliberately collapsed the six positions into only three bins of early, middle, and late. This reduced the number of distinguishable temporal patterns from 10 to 4 and the number of significant gene-pattern instances from 33 to 11. All 10 significant non-monotonic gene-pattern instances disappeared, and the interior structure of eight of nine non-monotonic biological-program dynamics was lost. The intermediate molecular states are therefore not merely a denser visualization of the trajectories; they preserve transient and stage-specific biological structure that is merged or lost at coarser temporal resolution. Intermediate molecular states recovered by CohortFM increase the effective biological resolution of a sparsely sampled longitudinal dataset. They separate molecular states that calendar visits collapse together and enable biological dynamics to be characterized at positions that are not accessible from the original three-visit representation alone.

#### 7.5.1 Biological Patterns Revealed by Higher-Resolution Progression

The previous analysis showed that the finer molecular progression scale increases biological resolution overall. Here we examine what specific biological information becomes visible at that resolution, focusing on concrete examples: programs whose temporal structure is missed by coarse visit-level sampling, whose dynamics differ across cohort-level trajectories, or whose signals concentrate at intermediate progression positions.

Two representative findings illustrate this added information. First, antiviral defense and interferon was broadly shared across trajectories but showed markedly different timing. Its strongest signal occurred at *s* = 0 in **T**_1_ and at *s* = 1 in **T**_2_. A calendar-only analysis pools these trajectories into a single D0–D3–D7 profile and therefore cannot recover this trajectory-dependent timing. Second, antigen presentation and lymphocyte immunity in **T**_5_ followed a late-dip pattern across the six molecular positions. The same biology appeared as an increasing pattern when samples were grouped only by calendar visit and as a decreasing pattern after the trajectory was reduced to three progression bins. Therefore, the higher-resolution representation exposes a transient structure that is not preserved by either coarse representation. These examples complement the resolution analysis by showing that the intermediate molecular states do not merely increase the number of detectable biological associations. They can change what can be concluded about *when* a biological program is active and *how* it evolves along a particular progression trajectory.

## 8 Generalization Across Datasets and Baselines

This section examines whether CohortFM generalizes beyond the COVID-19 cohort and whether the results reported earlier are reproducible across different diseases and sampling settings. The COVID-19 analysis was intentionally comprehensive to establish the biological capabilities of the framework in depth. Having established that foundation, the goal here is different. This section aims to test *generalization* through a more compact and standardized set of experiments across additional datasets. We therefore focus on a smaller set of biological evaluations that capture the main properties of the learned trajectories without repeating the full COVID-19 analysis for every cohort. Results for this experiment are provided in Section 8.1.

Beyond biological interpretation, CohortFM is ultimately a machine-learning model and should also generate accurate molecular states from sparse observations. We therefore evaluate its predictive capabilities and compare it with two state-of-the-art neural baselines, namely, scNODE, which is based on neural ODEs, and scFM, which also uses flow matching. These comparisons address two complementary questions: *i*) whether CohortFM is competitive with strong predictors, and *ii*) if its performance comes simply from using flow matching or from our particular design introduced for sparse longitudinal patient data. See Section 8.2 for our comparative results.

Finally, in Section 8.3, we examine the design of CohortFM itself through an ablation study. The training objective, from Equation 11 combines flow matching (*L*_*FM*_) with path-consistency (*L* _*path*_) and anchor supervision (*L*_*anchor*_), and we systematically remove or combine these components to understand their contribution. The results show that flow matching provides the essential learning signal, path consistency offers the strongest complementary benefit, and the complete objective provides consistently strong performance across heterogeneous datasets. The aforementioned three sections evaluate CohortFM from the perspectives of biological generalization, predictive accuracy, and the contribution of its core design choices.

### 8.1 Biological Generalization Across Disease Cohorts

We compared CohortFM with Truffle [Hasanaj et al., 2024] across COVID-19, sepsis, severe influenza, and tuberculosis. Truffle is a strong baseline for recovering recurrent routes through a graph of discrete molecular states. CohortFM addresses a related but different problem and it learns continuous patient-specific paths and subsequently identifies cohort-level trajectory archetypes. Our evaluation therefore does not ask whether one method is universally preferable. Instead, it asks which representation is more useful when the objective is to assign longitudinal patients to biologically informative cohort trajectories.

To compare the methods fairly, each sample was described by its trajectory and its stage. A stage is a discrete position along disease progression (such as an early, intermediate, or late molecular state). It describes where the patient is at the time of sampling. The trajectory represents the broader disease course that the patient follows across visits. Therefore, two patients may be at the same stage while following different trajectories. Only real patient samples were evaluated. CohortFM-generated intermediate states were excluded, and its continuous progression scale was converted into discrete stages to match Truffle. Each patient was evaluated by hiding their gene-expression data and estimating it from similar patients in the training set. The estimated expression was then compared with the patient’s actual measurements. Patients who could not be assigned to a trajectory were given the cohort-average estimate rather than being removed. This prevented methods with lower patient coverage from gaining an unfair advantage.

The first comparison measured whether trajectory identity added information beyond the local disease position or state, calculated as Δ*R*^2^ = *R*^2^_stage×state_ − *R*^2^_trajectory×stage_, where the reference was stage alone for CohortFM and state alone for Truffle.^10^ Results for this experiment are reported in Figure 11a. CohortFM trajectory identity increases held-out gene-expression *R*^2^ in all four diseases, with gains from 0.046 to 0.107. The corresponding Truffle changes were negative, ranging from −0.114 to − 0.190. These negative increments do not mean that Truffle removes information from the original data or that its state model is uninformative. Truffle states are obtained directly from the expression space and can provide a strong description of local molecular state. The negative increment instead means that compressing those states into matched cohort routes was less predictive of an unseen patient’s expression than retaining the state labels alone. This can occur when several informative states are merged into a common route, when route-specific groups become sparse, or when a patient cannot be matched reliably. Conversely, CohortFM is trained around patient-level paths, and its positive increments show that trajectory identity supplies information beyond progression position.

**Figure 11.**
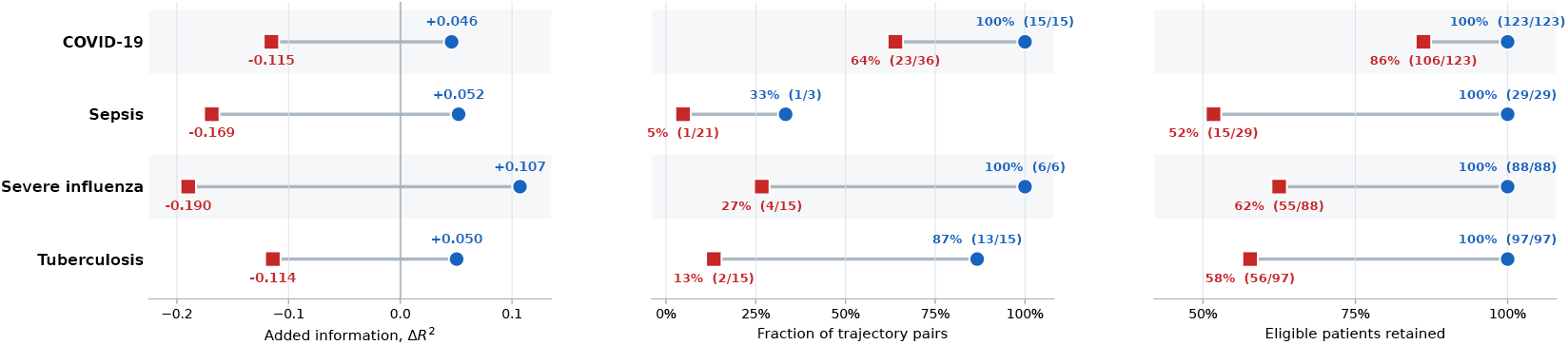
Cross-disease comparison of CohortFM and Truffle as cohort-trajectory abstractions. Each row represents one disease. **a**, Information contributed by trajectory identity, measured as the change in patient-held-out gene-expression *R*^2^ relative to each method’s non-trajectory representation. **b**, Fraction of proposed trajectory pairs that are biologically distinguishable using leave-one-patient-out classification of stage-residual co-expression-module profiles and 200 label permutations. **c**, Patient retention by both methods. Exact trajectory-pair and patient counts are shown beside each marker. 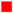represents Truffle and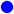 is CohortFM.

The second test is designed to examine whether different trajectories represented different biological patterns rather than simply different disease stages. The results are summarized in Figure 11b. We removed the effect of stage and then tested whether gene-expression patterns could correctly distinguish patients assigned to each pair of trajectories. Each patient was tested using information learned from the remaining patients. We also repeated the analysis after randomly shuffling the trajectory labels 200 times to determine whether the observed differences were greater than expected by chance. For CohortFM, 15 of 15 trajectory pairs were distinguishable in COVID-19, 1 of 3 in sepsis, 6 of 6 in severe influenza, and 13 of 15 in tuberculosis. For Truffle, the corresponding results were 23 of 36, 1 of 21, 4 of 15, and 2 of 15. Sepsis was challenging for both methods, but CohortFM produced a higher proportion of biologically distinct trajectories in all four datasets.

The third test measured how completely each method represented the patient cohort, as shown in Figure 11c. Among patients who could be evaluated by both methods, CohortFM retained all patients across four datasets. Truffle retained 86%, 52%, 62%, and 58%, for COVID-19, sepsis, influenza, tuberculosis, respectively. These differences reflect how the methods construct trajectories. Truffle assigns a patient only when their sequence of molecular states matches a recurrent route with sufficient overlap. This produces a focused description of common state transitions but leaves some patients without a complete trajectory–stage assignment. CohortFM first constructs a path for each patient and then assigns that path to a cohort-level trajectory, allowing it to retain all eligible patients. Our comparative analyses are not limited to the three tests reported in this section. See Appendix D for detailed results.

### 8.2 Interpolation and Forward Prediction Across Diseases

The utility of CohortFM ultimately depends on the accuracy of its predictions and reconstructions. We therefore evaluated its predictive performance across COVID-19, sepsis, severe influenza, and tuberculosis under two distinct settings. In the *interpolation* setting, an intermediate visit was withheld while observations on both sides of the missing state remained available. In the more stringent *forecasting* setting, the final visit was withheld and all methods received only observations preceding the target. Results for both settings are reported in Table 3.

**Table 3.** Interpolation and forward-prediction benchmarks across four longitudinal disease cohorts. Panel A evaluates reconstruction of a hidden internal visit. scFM and scNODE are not designed for interpolation at intermediate time points. Panels B and C evaluate true forward prediction, where all methods receive only observations preceding the target. Mean squared error (MSE) denotes the median across patients of each patient’s mean squared decoded-gene error, while sliced Wasserstein (SW) measures agreement between predicted and observed cohort-level gene-expression distributions (lower is better). Bold indicates the best result within a dataset and task, and underlining indicates the second-best.

| Dataset | Prediction task | CohortFM | Linear | scFM | scNODE |
| --- | --- | --- | --- | --- | --- |
| *A. Bracketed interpolation: patient-level gene MSE / gene SW* |  |  |  |  |  |
| COVID-19 | $D0, D7 \rightarrow D3$ | 0.988 / **0.357** | **0.880** / 0.371 | – | – |
| Sepsis | $D1-D3, D5 \rightarrow D4$ | 0.210 / 0.309 | **0.091** / **0.228** | – | – |
| Influenza | $T1, T3 \rightarrow T2$ | **0.242** / **0.313** | 0.242 / 0.316 | – | – |
| Tuberculosis | $W0, W1, W24 \rightarrow W4$ | **0.169** / 0.201 | 0.173 / **0.198** | – | – |
| *B. Forward prediction: patient-level gene MSE* |  |  |  |  |  |
| COVID-19 | $D0, D3 \rightarrow D7$ | **0.829** | 1.073 | 1.228 | 1.302 |
| Sepsis | $D1-D3 \rightarrow D4$ | **0.128** | <u>0.180</u> | 0.402 | 0.673 |
| Influenza | $T1, T2 \rightarrow T3$ | <u>0.383</u> | 0.773 | **0.367** | 0.442 |
| Tuberculosis | $W0, W1, W4 \rightarrow W24$ | **0.193** | <u>0.271</u> | 0.324 | 0.426 |
| *C. Forward prediction: gene-expression sliced Wasserstein (SW)* |  |  |  |  |  |
| COVID-19 | $D0, D3 \rightarrow D7$ | <u>0.324</u> | **0.311** | 0.455 | 0.515 |
| Sepsis | $D1-D3 \rightarrow D4$ | **0.264** | <u>0.293</u> | 0.396 | 0.554 |
| Influenza | $T1, T2 \rightarrow T3$ | **0.342** | <u>0.437</u> | 0.487 | 0.470 |
| Tuberculosis | $W0, W1, W4 \rightarrow W24$ | **0.165** | <u>0.228</u> | 0.291 | 0.297 |

The standard CohortFM formulation (Equation 4) requires the final state, denoted by *z*_1_, as an input. Because this state is precisely what must be predicted in the forecasting experiment, the standard endpoint conditioning cannot be used directly. We therefore retained the same CohortFM flow-matching backbone and replaced only the unavailable endpoint conditioning with information from the observed patient history. Specifically, the model uses the earliest and most recent observed visits, their molecular change, and the time to the target visit. Starting from the latest observed state, the same learned flow-matching dynamics are integrated forward to predict the patient’s future latent state. During training, future visits from training patients provide supervision, whereas the target visit remains completely hidden for test patients.

To provide a stronger evaluation, we included simple linear models as competitive baselines. Straight-line interpolation has access to the patient’s future endpoint and therefore represents a particularly strong baseline for reconstructing a missing intermediate visit. In contrast, the forecasting experiment enables a like-for-like comparison of CohortFM with linear extrapolation, scFM, and scNODE, since none of the methods has access to the future endpoint.

The interpolation results show that CohortFM remains competitive with a strong linear baseline when both endpoints are observed. It achieves the best distributional reconstruction in COVID-19 and influenza, matches linear interpolation closely in tuberculosis, and is less effective in the densely sampled sepsis setting, where a straight-line assumption is already strong. The advantage of CohortFM is clearer in forward prediction. Across the four diseases, CohortFM is consistently among the strongest methods at both the patient and cohort levels. It achieves the lowest patient-level MSE in COVID-19, sepsis, and tuberculosis, and the lowest sliced-Wasserstein distance in sepsis, influenza, and tuberculosis. Influenza is the only case where scFM yields a slightly lower patient-level MSE, while CohortFM still provides the best cohort-level distributional reconstruction. These results show that CohortFM provides a strong nonlinear predictor of molecular state transitions, supporting its use as the foundation for reconstructing and building continuous disease trajectories in our experiments.

### 8.3 Cross-Disease Ablation Supports the Training Design

CohortFM is trained with three complementary objectives of flow matching (FM), path consistency (Path), and anchor supervision. To understand the contribution of each objective, we performed a systematic ablation study across all four disease datasets. For each dataset, we evaluated all seven non-empty combinations of the three losses using three matched random seeds, while keeping the model architecture, patient splits, training budget, and all other settings unchanged. We compared all loss combinations using held-out hidden-visit prediction MSE, providing a common and consistent evaluation across configurations and datasets. Results are reported in Table 4.

**Table 4.** Cross-disease ablation of the three training objectives. Values show the mean *±* standard deviation of held-out hidden-visit prediction MSE across three seeds. The number in parentheses gives the rank within each dataset (1 = best). Bold marks the best result and underlining the second-best. The final column reports the average rank across all four datasets.

| Loss | COVID-19 | Sepsis | Influenza | Tuberculosis | Avg. rank |
| --- | --- | --- | --- | --- | --- |
| FM only | 2.0330 ± 0.0155 (3) | **1.1354 ± 0.0399 (1)** | 0.7938 ± 0.0128 (3) | 1.0753 ± 0.0034 (3) | 2.50 |
| Path only | 2.4715 ± 0.0149 (7) | 2.7089 ± 0.3574 (7) | 1.2917 ± 0.0615 (7) | 1.5014 ± 0.0180 (7) | 7.00 |
| Anchor only | 2.1112 ± 0.0072 (5) | 1.5125 ± 0.0719 (5) | 0.9394 ± 0.0092 (5) | 1.3204 ± 0.0236 (6) | 5.25 |
| FM + Path | **1.9401 ± 0.0697 (1)** | 1.1722 ± 0.0309 (3) | 0.7700 ± 0.0210 (2) | **1.0088 ± 0.0034 (1)** | **1.75** |
| FM + anchor | 2.0396 ± 0.0121 (4) | 1.1754 ± 0.0355 (4) | 0.8002 ± 0.0065 (4) | 1.0790 ± 0.0175 (4) | 4.00 |
| Path + anchor | 2.2563 ± 0.0160 (6) | 1.9879 ± 0.1866 (6) | 0.9901 ± 0.0089 (6) | 1.3014 ± 0.0481 (5) | 5.75 |
| All three | <u>2.0221 ± 0.0520 (2)</u> | <u>1.1678 ± 0.0337 (2)</u> | **0.7654 ± 0.0165 (1)** | <u>1.0134 ± 0.0765 (2)</u> | **1.75** |

Several consistent patterns emerge from Table 4. First, FM is the essential component of CohortFM. Every configuration that excludes the FM objective performs substantially worse. The auxiliary losses therefore cannot replace learning the underlying patient-specific vector field.

Second, Path provides the clearest complementary benefit. Adding the path-consistency loss to FM improves hidden-visit prediction over FM alone. FM + Path is the best configuration for COVID-19 and tuberculosis and the second-best for influenza. This result supports the use of observed intermediate visits as additional supervision.

FM + anchor does not outperform FM alone, which indicates the anchor objective should be viewed as a secondary objective/regularizer. However, when incorporated at its relatively small weight into the full objective, performance remains consistently strong. The complete three-loss model ranks among the top two configurations in every dataset and achieves the best result for influenza. Its average rank of 1.75 across diseases ties FM + Path for the strongest overall performance.

These results suggest that each loss plays a distinct role and that no single combination dominates every dataset. Although FM + Path and the full objective have the same average rank, we retain all three losses as the default configuration because the full model is consistently among the top two across all datasets while also achieving the best performance on influenza. The anchor term provides an additional constraint on the learned trajectories at a small weight, without sacrificing overall predictive performance. We therefore favor the three-loss objective as the more complete and robust default, while recognizing that FM + Path performs essentially on par.

## 9 Conclusion and Future Work

We introduced CohortFM, a framework for transforming sparse longitudinal molecular observations into continuous patient-specific trajectories and a compact set of cohort-level progression patterns. The framework separates molecular progression from calendar time, learns continuous transitions between observed molecular states, places patients on a shared progression coordinate, and maps the resulting trajectories back to real patient samples for biological interpretation. Across COVID-19, sepsis, severe influenza, and tuberculosis, CohortFM recovered biologically informative progression structure and generalized across substantially different cohort sizes and sampling schedules. Its interpolation and forward-prediction performance further support the learned trajectories as useful representations of longitudinal molecular dynamics.

An important direction for future work is to further improve the machine-learning component of the framework. This setting presents an inherent challenge. Expressive models such as flow matching can capture complex nonlinear dynamics, but they typically benefit from larger training datasets, while longitudinal molecular datasets are often small and sparsely sampled. Simpler non-machine-learning approaches require less data, but may struggle to represent the same level of heterogeneity and nonlinear progression. Bridging this gap remains an important research opportunity. Future work could therefore focus on more data-efficient learning, stronger regularization, transfer learning across related cohorts, and methods that better exploit shared structure while remaining robust in low-data regimes.

A second direction is to move beyond a single molecular modality. Transcriptomics provides only one view of disease progression, and it would be particularly valuable to study cohorts in which the same patients have longitudinal measurements across multiple modalities, such as proteomics, metabolomics, clinical variables, treatment history, digital-health data and other modalities. Extending CohortFM to such multi-modal trajectories could provide a richer representation of progression and help determine whether different modalities evolve synchronously or capture complementary aspects of disease.

Future studies should also evaluate transfer across independent cohorts and investigate whether trajectories learned in one population can be used to position or forecast patients from another. Prospective longitudinal datasets with denser sampling would provide particularly valuable validation, since they would allow intermediate states predicted from artificially sparse observations to be compared directly with measurements that were actually collected.

Finally, the broader problem considered here extends beyond molecular disease modeling. Many longitudinal datasets consist of discrete and irregular observations of processes that evolve continuously. CohortFM provides one approach for using population-level information to reconstruct the missing structure between those observations while preserving individual trajectories. Exploring this principle in other sparse longitudinal domains represents a natural direction for future work.

## Supporting information

dataset preparation, model training details, and complementary results

## Footnotes

2 This technical report is a work in progress. Supporting code and analyses are available upon request.

3 AI was used to assist with paraphrasing and improving the clarity of the manuscript. All scientific questions, study design, experiments, analyses, and interpretations were developed, conducted, and supervised by the authors. AI was used solely to accelerate the writing process.

4 https://www.ncbi.nlm.nih.gov/geo/query/acc.cgi?acc=GSE212041

5 GSE54514 is reported as comprising 35 patients, although its sample annotations contain 36 unique patient identifiers.

6 The 3*d* terms are the current state **z**_*τ*_, source **z**_0_, and target **z**_1_. The 14 scalar features are *τ*, its 8 sinusoidal encodings, the pseudotimes *s*_*τ*_, *s*_0_, *s*_1_, the pseudotime displacement !*s*, and the normalized calendar-time displacement 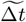.

7 The model evaluates several candidate values and selects the one that minimizes the reconstruction loss.

8 For each trajectory–position cloud containing at least three samples, expression was compared with 87 pooled COVID-negative controls using Welch’s *t*-test on log_2_(TPM + 1). Genes were tested if TPM ≥1 in at least 10% of the cloud-plus-control samples, with Benjamini–Hochberg correction applied within each cloud; significance required adjusted *P* < 0.05 and |log_2_ FC| ≥ 0.5. Up- and downregulated genes were analyzed separately, using at most the 200 genes with largest absolute fold change per direction and requiring at least 10 unique symbols. GO enrichment used Enrichr GO Biological Process 2023 for Human with its default background; terms with Enrichr-adjusted *P* < 0.05 were retained, with no additional correction across cloud–direction queries.

9 For each trajectory, gene-expression profiles across the six normalized positions were standardized and matched to predefined temporal patterns. Genes were retained when their expression amplitude was at least 0.5 log_2_ units and their pattern correlation was *r* ≥ 0.80. Pattern enrichment was tested against all 6! = 720 permutations of the position order, with Benjamini–Hochberg correction across trajectory–pattern tests; significance required *q <* 0.05 and at least 30 genes. Up to 300 best-fitting genes per significant pattern were submitted to Enrichr GO Biological Process 2023 using the 2,000-gene expression panel as background, retaining GO terms with adjusted *P <* 0.05.

10 *R*^2^ measures how well the labels predict gene-expression differences in patients who were not used to build the prediction; ×denotes a joint trajectory–stage label rather than multiplication.

