## Supplementary material for "From Sparse Visits to Continuous Molecular Trajectories with Flow Matching": dataset preparation, model training details, and complementary results

### A Dataset Preprocessing and Quality Control

This section provides details on the preprocessing and quality-control procedures applied to the three supplementary longitudinal transcriptomic datasets used in our experiments.

**Sepsis.** The sepsis dataset (GSE54514),[11] contains Illumina HumanHT-12 v3 whole-blood microarray measurements from 36 patients admitted to the intensive care unit with sepsis and 18 healthy controls. Whole-blood samples were collected daily for up to five days following admission. Probes detected at $p < 0.05$ in at least $1\%$ of arrays were retained. Expression intensities were $\log_2$-transformed and quantile-normalized jointly across all 163 arrays. Probes were subsequently mapped to protein-coding genes. For cases where multiple probes mapped to the same gene, the probe with the highest mean expression was selected. Genes were then ranked according to their IQR across the sepsis samples, and the 2,000 most variable genes were retained.

**Severe Influenza.** The severe-influenza dataset (GSE111368),[12] contains Illumina HumanHT-12 v4 whole-blood microarray measurements from 109 patients with influenza and 130 healthy controls. Longitudinal samples were collected at three clinically relevant stages: T1 (recruitment/enrollment) that corresponds to the day of hospital admission, T2 (acute follow-up) that was collected approximately two days after T1, and T3 that belongs to the convalescent or recovery stage. Probes were required to have a detection score of at least 0.95 in four or more arrays. Background-corrected intensities were $\log_2$-transformed, quantile-normalized, and corrected for BeadChip-associated effects using ComBat. Probes were mapped to protein-coding Ensembl genes. The 2,000 genes with the largest IQR were selected for downstream analysis.

**Tuberculosis.** The final dataset was the longitudinal tuberculosis (TB) cohort (GSE89403).[13] Samples were collected at four principal treatment stages: Week 0, corresponding to the initiation of TB treatment; Week 1, one week after treatment initiation; Week 4, four weeks after treatment initiation; and Week 24, approximately six months after the start of treatment. The dataset contains longitudinal whole-blood RNA-seq measurements collected during treatment. The original 914 sequencing records were collapsed into biological samples because a substantial number of samples contained technical replicates. Only patients with evaluable treatment outcomes were retained. The analysis was restricted to protein-coding genes, duplicate gene symbols were collapsed by retaining the row with the highest mean expression, and genes were required to have a CPM greater than $0.25$ in at least $1\%$ of samples. Counts were TMM-normalized and transformed using $\log_2(\mathrm{CPM}+1)$. Sample-level GC-content bias was corrected using 50 GC-content bins. The resulting dataset contained 367 samples from 98 patients. As with the other datasets, the 2,000 genes with the highest IQR were retained as the final input representation.

### B Regularization of the Cohort-Level Mixture Model

The mixture model is trained primarily to reconstruct each patient trajectory as a probability-weighted combination of the learned cohort-level trajectories. Reconstruction alone, however, may lead to trajectories that are irregular, redundant, poorly utilized, or associated with overly diffuse patient assignments. We therefore add four regularization terms.

**Trajectory smoothness.** To encourage each cohort-level trajectory to evolve smoothly along the normalized progression axis, we penalize both first- and second-order changes:

$$\mathcal{L}_{\text{smooth}} = \mathbb{E}_{\mathrm{t},m}\left[\|\mathbf{T}_{\mathrm{t},m+1} - \mathbf{T}_{\mathrm{t},m}\|_2^2 + \|\mathbf{T}_{\mathrm{t},m+2} - 2\mathbf{T}_{\mathrm{t},m+1} + \mathbf{T}_{\mathrm{t},m}\|_2^2\right] \tag{14}$$

**Balanced trajectory utilization.** To discourage the model from concentrating most patients on only a few cohort-level trajectories, we define the average assignment probability

$$: \bar{\pi}_{\mathrm{t}} = \mathbb{E}_p\left[\pi_{p,\mathrm{t}}\right], \tag{15}$$

[11] https://www.ncbi.nlm.nih.gov/geo/query/acc.cgi?acc=GSE54514

[12] https://www.ncbi.nlm.nih.gov/geo/query/acc.cgi?acc=GSE111368

[13] https://www.ncbi.nlm.nih.gov/geo/query/acc.cgi?acc=GSE89403

and penalize strong deviations from balanced utilization:

$$\mathcal{L}_{\text{balance}} = \mathbb{E}_{\text{t}} \left[ \left( \bar{\pi}_{\text{t}} - \frac{1}{\mathcal{T}} \right)^2 \right] \tag{16}$$

**Assignment confidence.** We use an entropy penalty to discourage excessively diffuse patient assignments while retaining the soft-mixture formulation:

$$\mathcal{L}_{\text{conf}} = -\mathbb{E}_p \left[ \sum_{\text{t}=1}^{\mathcal{T}} \pi_{p,\text{t}} \log \pi_{p,\text{t}} \right] \tag{17}$$

**Trajectory diversity.** To prevent different cohort-level trajectories from collapsing onto nearly identical patterns, we apply a margin-based penalty:

$$\mathcal{L}_{\text{div}} = \mathbb{E}_{(i,j)\in\mathcal{Q}} \left[ \max\left(0,\ \delta - \|\mathbf{T}_i - \mathbf{T}_j\|_F\right)^2 \right], \tag{18}$$

where $\mathcal{Q}$ is the set of trajectory pairs and $\delta$ is a diversity margin. In our implementation, $\delta$ is set to one quarter of the median pairwise distance between patient trajectories in the training data.

The complete mixture-model objective is a weighted combination of all these components.

### C Trajectory-Specific Composition of Calendar Visits Along Molecular Progression

Figure 12 provides the trajectory-specific version of the pooled analysis presented in Section 7.1. For each CohortFM archetype, $\mathbf{T}1$–$\mathbf{T}6$, we show the proportion of D0, D3, and D7 observations assigned to each normalized molecular-progression position $s \in \{0, 0.2, 0.4, 0.6, 0.8, 1.0\}$. The number above each bar indicates the number of real patient samples contributing to that trajectory-position combination. Across trajectories, the expected broad temporal organization is preserved: D0 samples are generally concentrated toward earlier molecular positions, whereas D7 samples become increasingly prevalent toward later positions. However, the detailed composition differs substantially between archetypes.

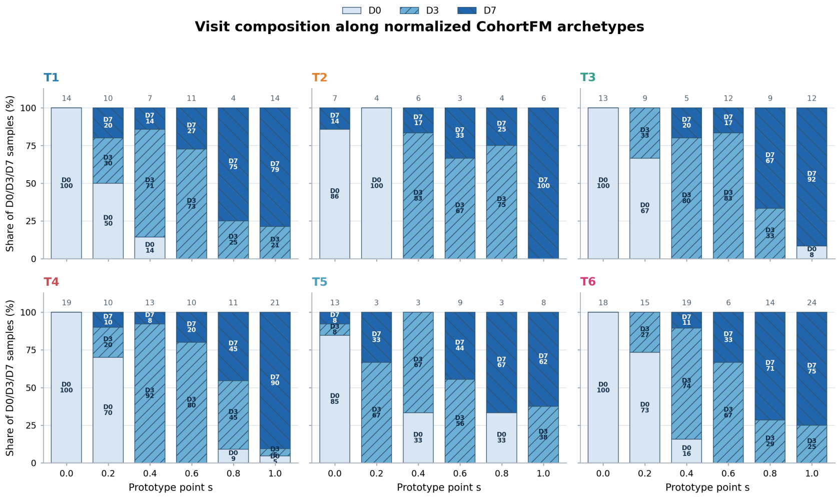


Figure 12: Trajectory-specific composition of clinical visits along the normalized molecular-progression axis. For each CohortFM archetype ($\mathbf{T}_1$–$\mathbf{T}_6$), stacked bars show the proportions of D0, D3, and D7 samples assigned to each normalized progression position. Numbers above the bars indicate the number of patient samples contributing to each position.

### D Cross-Disease Comparison

#### D.1 COVID-19

Although the Truffle trajectory–stage abstraction achieves a slightly higher absolute held-out gene-expression $R^2$ than CohortFM in COVID-19 ($0.186$ versus $0.159$), the two quantities capture different aspects of the representation. Much of Truffle's absolute predictive performance is already contained in its underlying state representation. Adding trajectory identity reduces held-out $R^2$ by $0.115$, whereas adding CohortFM trajectory identity to its stage representation increases $R^2$ by $0.046$. Thus, the COVID-19 results illustrate the distinction between representing a local molecular state and learning a cohort-level trajectory that contributes information beyond that state.

The coverage analysis further highlights a methodological difference between the approaches. CohortFM first reconstructs a patient-specific path and then maps that path to a cohort-level trajectory, retaining all 123 patients in the common three-visit population. Truffle retains 106 of these patients under the matched trajectory criterion, although it can additionally represent some patients with shorter longitudinal profiles. The retention comparison should therefore be interpreted as the robustness of the resulting cohort-level trajectory abstraction under the common evaluation setting rather than as a general limitation on Truffle's ability to process shorter trajectories.

Figure 13 provides a more detailed view of the biological structure captured by the two representations. The trajectory–stage profiles contain variation across multiple biological programs, including antiviral and inflammatory responses, lymphocyte activation, antigen presentation, mitochondrial organization, translation, and cell-cycle processes. These profiles complement the aggregate comparisons in Section 8 by showing how molecular programs are organized across individual trajectory–stage states.

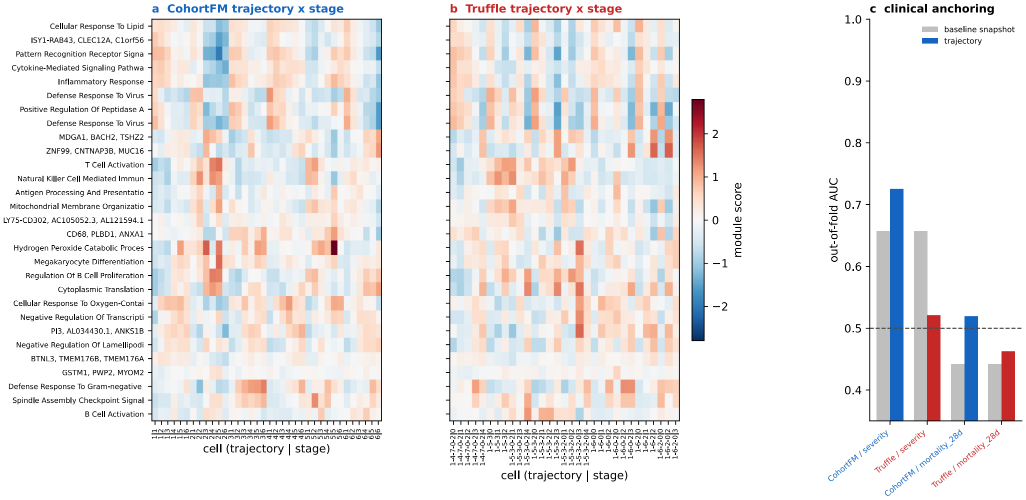


Figure 13: Biological profiles and clinical anchoring of the COVID-19 trajectory representations. Heatmaps show biological-module activity across trajectory–stage states for CohortFM and Truffle. The final panel evaluates the association of the learned trajectory representations with COVID-19 severity and 28-day mortality using out-of-fold prediction.

As a secondary clinical check, we asked whether the learned trajectory representation contains information about patient outcomes that was not used to train either method. For COVID-19 severity, a baseline snapshot model achieves an out-of-fold AUC of $0.657$. Incorporating the CohortFM trajectory increases this to $0.725$, whereas the Truffle trajectory gives an AUC of $0.521$. For 28-day mortality, CohortFM increases AUC from $0.442$ to $0.519$, compared with $0.463$ for Truffle. Because mortality events are limited, this latter result should be interpreted cautiously. We therefore treat the clinical analyses as supporting evidence rather than part of the primary trajectory comparison.

#### D.2 Sepsis

In sepsis, the CohortFM trajectory–stage abstraction retains more held-out gene-expression variation than Truffle ($R^2 = 0.089$ versus $0.068$). Decomposing the representations again shows that trajectory identity contributes differently in the two approaches. Adding CohortFM trajectory identity to the stage representation increases held-out $R^2$ by $0.052$, whereas adding Truffle trajectory identity reduces $R^2$ by $0.169$ relative to the corresponding Truffle state representation. This suggests that, even in this relatively short and densely sampled cohort, the CohortFM trajectory contains information beyond the patient's local molecular state. The coverage difference is also substantial. CohortFM retains all 29 patients in the common evaluation population, whereas Truffle retains 15 of 29 patients under the matched trajectory criterion. As in COVID-19, this comparison reflects the robustness of the resulting cohort-level trajectory abstraction under the common evaluation setting rather than a general inability of Truffle to represent individual molecular states.

Figure 14 provides a more detailed view of the biological structure represented across trajectory–stage states. The profiles span processes related to translation and peptide biosynthesis, antigen-receptor signaling, innate and antimicrobial responses, regulation of cell death, endocytosis, heme metabolism, and inflammatory signaling. The structure is less strongly separated than in COVID-19, which is consistent with the smaller cohort and the relatively dense sampling of sepsis over a short time window. As a secondary clinical check, we evaluated whether the trajectory representations were associated with mortality. Both methods produced only modest out-of-fold discrimination, with AUCs slightly above chance and no clear advantage for CohortFM.

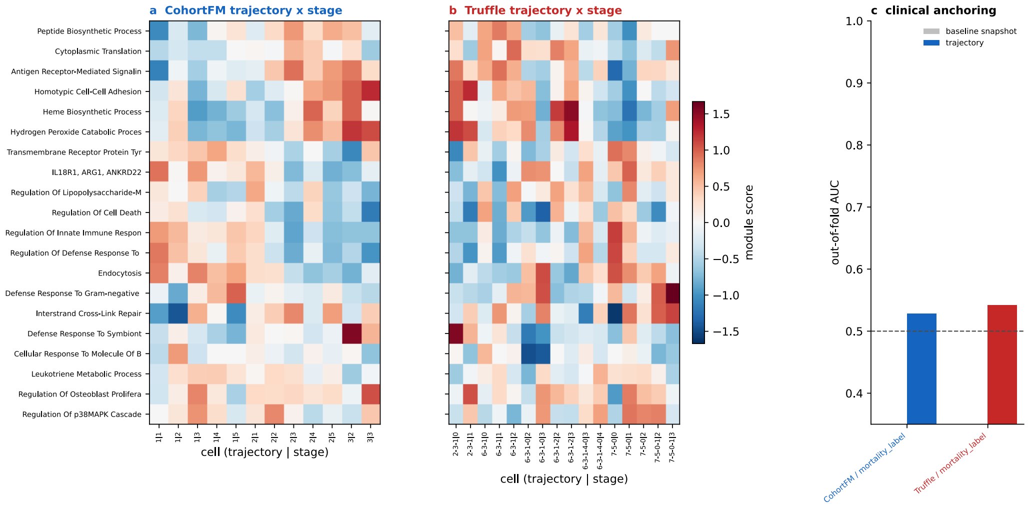


Figure 14: Biological profiles and clinical anchoring of the sepsis trajectory representations. Heatmaps show biological-module activity across trajectory–stage states for CohortFM and Truffle. The final panel evaluates the association of the learned trajectory representations with mortality using out-of-fold prediction.

#### D.3 Severe Influenza

In severe influenza, decomposition of the trajectory representations shows a particularly clear difference between the two approaches. Adding CohortFM trajectory identity to the stage representation increases held-out gene-expression $R^2$ by $0.107$, whereas adding Truffle trajectory identity reduces $R^2$ by $0.190$ relative to the corresponding Truffle state representation. Thus, trajectory identity in CohortFM captures substantial molecular information beyond the patient's local progression state.

The coverage analysis shows a similar difference at the cohort level. CohortFM retains all 88 patients in the common evaluation population, whereas Truffle retains 55 of 88 patients under the matched trajectory criterion. As in the other datasets, this comparison reflects the completeness of the resulting cohort-level trajectory abstraction under the common evaluation setting rather than the ability of Truffle to represent individual molecular states.

Figure 15 provides a more detailed view of the molecular programs represented across trajectory–stage states. The profiles include inflammatory and antiviral responses, interleukin signaling, pattern-recognition pathways, B-cell and immunoglobulin-mediated immunity, platelet activation, oxygen transport, and translation-related processes. These profiles complement the aggregate results provided in the main body of the report by showing how biological programs vary across individual progression paths and stages.

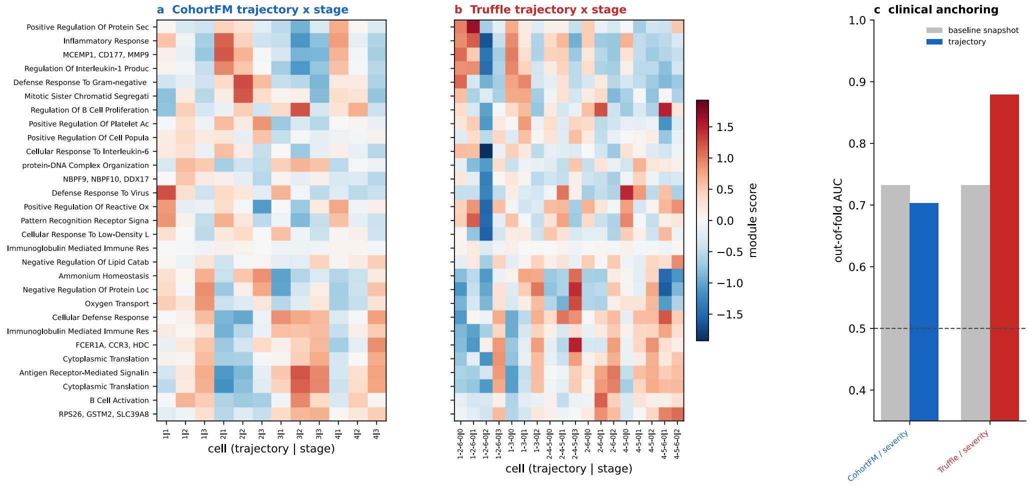


Figure 15: Biological profiles and clinical anchoring of the severe-influenza trajectory representations. Heatmaps show biological-module activity across trajectory–stage states for CohortFM and Truffle. The final panel evaluates the association of the learned trajectory representations with disease severity using out-of-fold prediction.

As a secondary clinical check, we evaluated whether the trajectory representations improved prediction of disease severity. In contrast to the molecular analyses above, the CohortFM trajectory does not improve over the baseline snapshot representation, whereas the Truffle trajectory produces a substantially higher out-of-fold severity AUC. This result highlights that capturing biologically informative longitudinal structure and predicting a specific clinical outcome are related but distinct objectives. Because severity labels were not used to train either trajectory model, we treat this analysis as an external clinical check rather than part of the primary trajectory comparison.

##### D.4 Tuberculosis

Tuberculosis provides a particularly challenging setting because observations span a much longer time scale, including a large gap between the early and late treatment measurements. CohortFM retains more held-out gene-expression variation than the Truffle trajectory–stage abstraction ($R^2 = 0.151$ versus $0.045$). Decomposing the representations shows that adding CohortFM trajectory identity increases held-out $R^2$ by $0.050$, whereas adding Truffle trajectory identity reduces it by $0.114$ relative to the corresponding Truffle state representation. Thus, even across widely separated observations, CohortFM trajectory identity contributes molecular information beyond the patient's local progression state.

The coverage analysis shows a substantial difference between the resulting cohort-level abstractions. CohortFM retains all 97 patients in the common evaluation population, whereas Truffle retains 56 of 97 patients under the matched trajectory criterion. As in the other cohorts, this comparison reflects the completeness of the trajectory abstraction under the common evaluation setting rather than a general limitation on Truffle's ability to represent individual molecular states.

Figure 16 provides a more detailed view of the biological structure represented across trajectory–stage states. The profiles include B-cell and innate immune activity, macrophage activation, antigen processing and presentation, antiviral and antibacterial responses, cellular respiration, translation, endocytosis, platelet activation, and oxidative-stress-related processes. These profiles complement the aggregate results in Section 8 by showing how molecular programs vary across individual progression paths and stages.

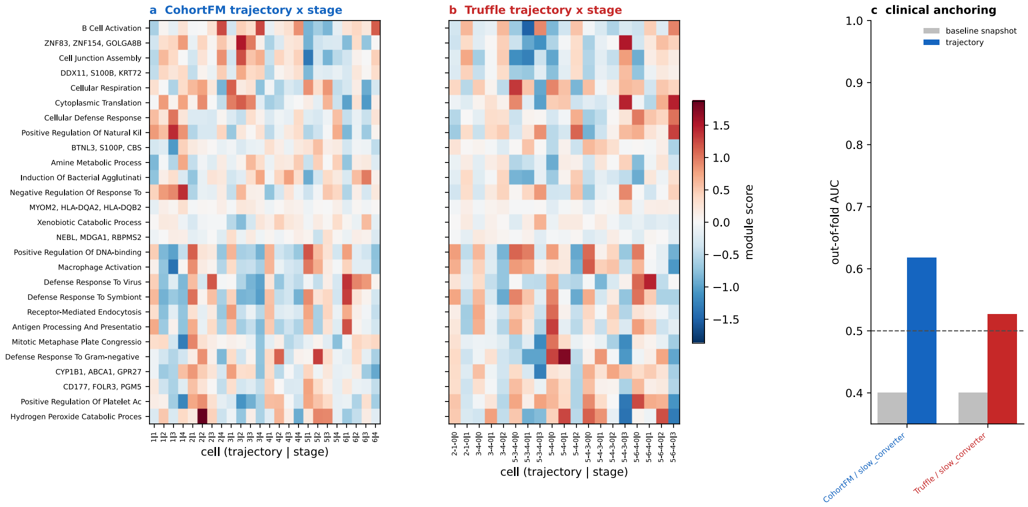


Figure 16: Biological profiles and clinical anchoring of the tuberculosis trajectory representations. Heatmaps show biological-module activity across trajectory–stage states for CohortFM and Truffle. The final panel evaluates the association of the learned trajectory representations with slow treatment conversion using out-of-fold prediction.

As a secondary clinical check, we evaluated whether the trajectory representations were associated with slow treatment conversion. The baseline snapshot representation provides little discriminative information, whereas both trajectory representations improve out-of-fold prediction. The improvement is larger for CohortFM than for Truffle, providing additional evidence that the learned longitudinal representation captures clinically relevant variation not evident from the baseline measurement alone. As with the other clinical analyses, this outcome was not used to train the trajectory models, and we therefore treat the result as an external check rather than part of the primary trajectory comparison.
